# Single molecule structure sequencing reveals RNA structural dependencies, breathing and ensembles

**DOI:** 10.1101/2020.05.18.101402

**Authors:** Teshome Tilahun Bizuayehu, Kornel Labun, Martin Jakubec, Kirill Jefimov, Adnan Niazi, Eivind Valen

**Affiliations:** Computational Biology Unit, Department of Informatics, University of Bergen, Norway; Sars International Center for Marine Molecular Biology, University of Bergen, Bergen 5008, Norway; Department of Chemistry, University of Tromsø, Norway

## Abstract

RNA molecules can form secondary and tertiary structures that can regulate their localization and function. Using enzymatic or chemical probing together with high-throughput sequencing, secondary structure can be mapped across the entire transcriptome. However, a limiting factor is that only population averages can be obtained since each read is an independent measurement. Although long-read sequencing has recently been used to determine RNA structure, these methods still used aggregate signals across the strands to detect structure. Averaging across the population also means that only limited information about structural heterogeneity across molecules or dependencies within each molecule can be obtained. Here, we present Single-Molecule Structure sequencing (SMS-seq) that combines structural probing with native RNA sequencing to provide non-amplified, structural profiles of individual molecules with novel analysis methods. Our new approach using mutual information enabled single molecule structural interrogation. Each RNA is probed at numerous bases enabling the discovery of dependencies and heterogeneity of structural features. We also show that SMS-seq can capture tertiary interactions, dynamics of riboswitch ligand binding, and mRNA structural features.

## Introduction

Chemical probing combined with sequencing is broadly used for RNA structural profiling (1). Most of these experiments use chemicals that modify accessible non-paired bases that result in reverse transcription termination and/or misincorporation of bases during cDNA synthesis (2–5). These sites can then be captured through sequencing the cDNA, resulting in an average reactivity score for each base. Computational tools based on mutational profiling have been developed to determine single molecule RNA structure determination (6, 7). While these methods have been instrumental in mapping structures across the transcriptome, they are susceptible to multiple biases (8). First, the synthesis of cDNA gives rise to biases through the processivity of reverse transcriptase, sequence biases during the random priming used to initiate the RT, and PCR amplification. Second, these methods are further constrained by the read length of short-read sequencers that prohibit full-length structural profiling of larger molecules such as messenger RNAs and ribosomal RNAs. Third, most of these methods only capture a single or very few modified bases per sequenced read. Finally, since most of these methods rely on amplification, the data obtained represents the average reactivity at a position across all molecules in the sample. These approaches are therefore limited in their ability to capture information about isoform-specific structures and structural heterogeneity.

Direct sequencing of long native RNA molecules after chemical probing can reveal insights into the complete structural profiles of longer RNA molecules (9, 10). This could lead to better prediction algorithms and, in turn, more accurate interpretations of RNA structures and functions. Here, we developed a novel method based on nanopore sequencing technology that directly detects the chemical adducts from structural probing on both short and long RNA molecules without the need for reverse transcription and PCR. Unlike two recent nanopore-based structure methods, which only consider average nucleotide accessibility and consensus structure (9, 10), our single-molecule approach probes multiple bases on individual molecules and describes their co-occurrence. We demonstrate this method on known structures and show that it can open the way for studying structural co-dependence of bases and conformational heterogeneity of RNA structures. We analyze the dynamics of riboswitches revealing conformational changes upon ligand binding, short- and long-range interactions with cooperative structural domains, and tertiary RNA interactions that have previously been described by crystal structures. In summary, we have developed a succinct structure probing approach that can provide a dynamic single-molecule view of RNA structure.

## Materials and methods

### Sample preparation

*In vitro* synthesis of RNA. Custom sense and antisense hairpin oligos with T7 promoter sequence were purchased from Sigma (Sigma-Aldrich) and *in vitro* transcribed using T7 RNA polymerase (NEB). *F. nucleatum* FMN riboswitch RNA was synthesized *in vitro* using T7 RNA polymerase from a pUT7 vector that carries 112-nucleotides FMN sensing domain with hammerhead ribozyme at the end to obtain identical 3′ ends (11). Coenzyme thiamine pyrophosphate binding riboswitch (TPP) RNA was synthesized *in vitro* using T7 RNA polymerase. TPP template was prepared using sense and antisense TPP oligos with T7 promoter sequence (**Supplementary Table 1**) purchased from Sigma (Sigma-Aldrich) that was assembled using PCR, cloned to pCR 2.1-TOPO vector (Invitrogen), and digested using EcoR I (NEB). Similarly, k-mer sequences were assembled using PCR from oligos purchased from Sigma.

Cell culture. Yeast (*Saccharomyces cerevisiae*) strain Y187 was grown in YPD medium at 30 °C. Saturated cultures were diluted to OD600 of ∼0.1 and grown to a final OD600 of 0.7 followed by RNA extraction using hot phenol-chloroform method. Poly(A) and non-poly(A) separation was performed using Poly(A)Purist MAG Kit (Invitrogen) following manufacturer protocol. The remaining non-poly(A) RNA was precipitated overnight at -80 °C following the RNA ethanol precipitation protocol.

Diethyl pyrocarbonate (DEPC) treatment of RNA. All structural probing was carried out using DEPC (Sigma), which carbethoxylates unpaired adenosine at N-6 or N-7 by opening the imidazole ring (12) (**Fig 1A**). 1μg RNA was denatured at 90 °C for 3 minutes, cooled on ice, and renatured in 200 mM Hepes pH 7.8, 100 mM KCl, 10 mM MgCl2 for 20 minutes at 20 °C. For (+)DEPC, RNA was treated with 10% (final volume) DEPC for 45 min at 20 °C. DEPC reaction was performed in the absence and presence of six-fold molar excess of riboflavin 5′-monophosphate and thiamine pyrophosphate (Sigma-Aldrich) for FMN and TPP riboswitch, respectively. The reactions were immediately purified using RNA Clean & Concentrator (Zymo Research) or RNA XP bead (Beckman Coulter).

**Fig. 1.**
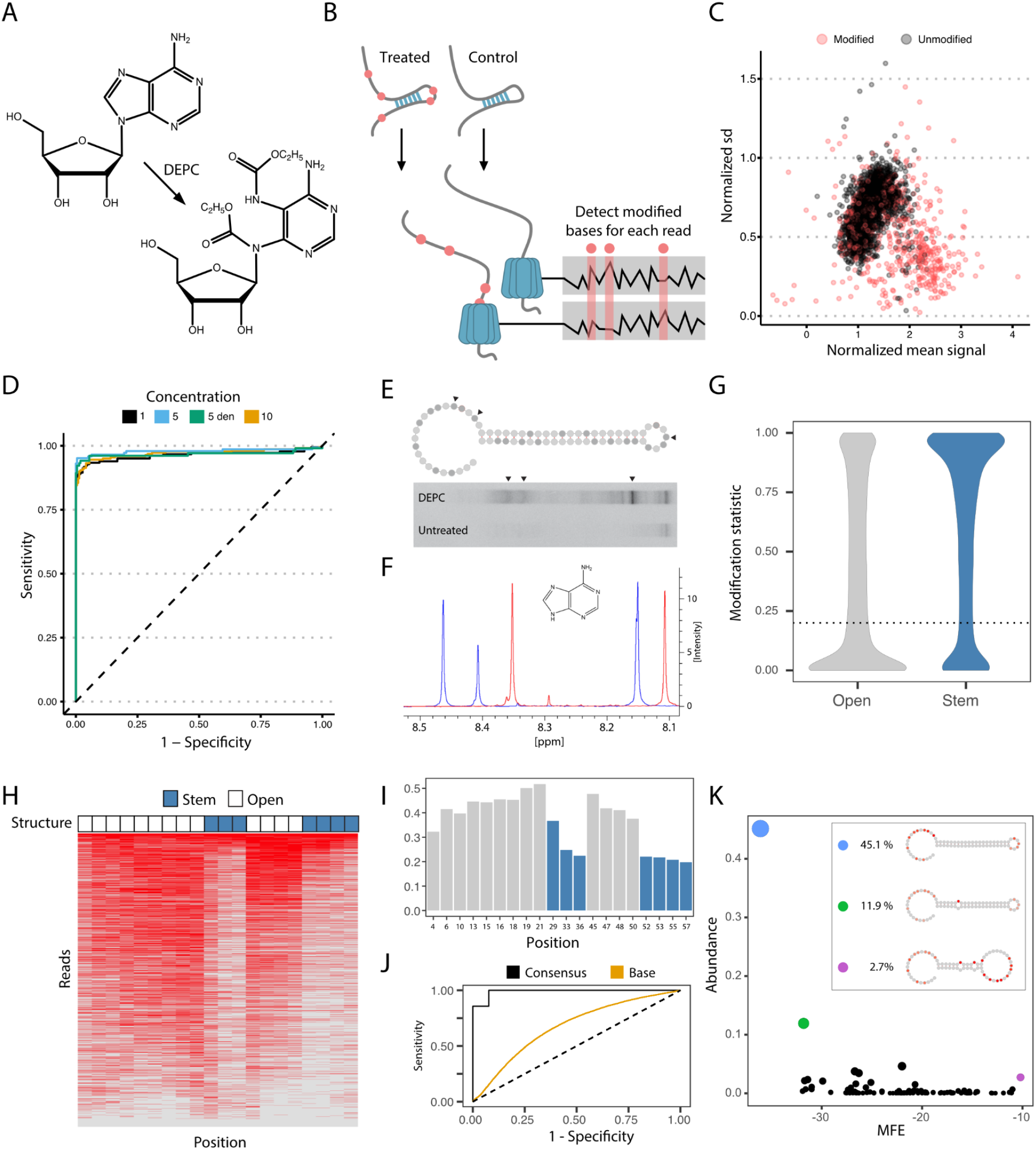
Oxford nanopore direct RNA structure profiling of a hairpin RNA. **A**) Molecular structure of Adenosine modification by DEPC that modifies structurally unconstrained adenosine by opening the imidazole ring. **B**) SMS-seq workflow. Structures are probed with DEPC and the current signal of each nucleotide that passes through the pore is recorded. Modified sites (red) are detected by comparing the current signal from treated samples to the control samples. **C**) Normalized current mean and standard deviation for a nucleotide at a single-stranded region of RNA modified with DEPC (red) and unmodified (gray). **D**) ROC curves for unmodified, modified, and denatured 5-mer RNA sequences. The 5-mer RNAs are treated with 1 % (black), 5 % (blue), 10 % (orange) DEPC as well as denatured RNA treated with 5 % DEPC (green). **E**) Denaturing 15% polyacrylamide gel electrophoresis of 5′-end radiolabeled RNA fragments generated from DEPC treated hairpin RNA (top) and untreated hairpin RNA (bottom), which were treated with borohydride and aniline. Arrows mark positions that were likely to be identified by SMS-seq as modified. **F**) Proton NMR spectra of non-treated (blue) and DEPC treated (red) for hairpin RNA. Singlet of protons on positions 8 and 2 on the purine ring could be quantified. **G**) Tombo modification statistics for adenosine bases. Bases are called modified (open) when their statistic is below 0.2 (dotted line) corresponding to an FDR of 15%. **H**) Heatmap showing 3093 full-length reads (y-axis) with modified (red) and unmodified (gray) nucleotides at each position (x-axis) for each read. Reads are ordered by the overall modification rate. The legend above (Structure) shows which nucleotides are accessible (white) and inaccessible (blue) in the hairpin structure. **I**) Modification frequency per base over the hairpin structure. Position 29 is uncharacteristic of a stem and is called as modified to a greater degree. This high modification rate could be due to RNA conformational heterogeneity, RNA breathing, or a problematic k-mer. **J**) ROC curve for the hairpin at the consensus level (black) and individual bases (orange). A random model is shown with the dotted line. **K**) Groups of reads folding to the same structure are shown by abundance (y-axis) and minimum folding energy (x-axis). The inset contains selected structures with modified bases colored (red circles).

### Oxford nanopore sequencing and mapping

Library preparation: Sequencing libraries were prepared following the direct RNA sequencing protocol (Oxford Nanopore Technologies, ONT) by replacing the 3′ adapters (RTA) with custom-made barcodes for multiplexing (**Supplementary Table 1**) (https://github.com/hyeshik/poreplex). After renaturation and treatment with DEPC, *in vitro* synthesized and yeast non-poly(A) RNA were polyadenylated using E.coli polyA polymerase (NEB) at 37 °C for 15 min. To obtain full-length reads and a molecule long enough to be processed, the hairpin and riboswitches were ligated to the 3′ ends of a longer RNA molecule (**Supplementary Table 1**). RTA adapter-ligated 500 ng RNAs were reverse-transcribed using superscript III. Different barcodes were used for the (+)DEPC and (−)DEPC libraries (**Supplementary Table 1**). The first-strand complementary DNAs were pooled and a motor protein ligated, which were subjected to Oxford Nanopore sequencing using MinION Flow Cell (R9.4) and run for different durations until we obtained what we deemed to be a sufficient number of reads.

Read mapping: Reads were demultiplexed using poreplex (v 0.5)(https://github.com/hyeshik/poreplex) and resquiggled using Tombo v1.5 (ONT)(13) by mapping to their respective references. Yeast reference transcriptome was obtained from Ensembl (version R64-1-1). Reads were mapped by allowing mismatches, insertions, and deletions to account for nanopore sequencing errors using minimap2 (v 2.9)(14).

### Detection of DEPC modification

Chemical probing of adenine residues with DEPC. Several studies indicate that DEPC can be used for detecting single-stranded adenosine in RNA molecules (15–20). Chemical reactivity is generally indicative of an absence of base stacking. Lack of reactivity, however, is more difficult to interpret since it could result not only from base pairing and tertiary interactions but also from the presence of stabilizing divalent cations (16).

End labeling: The ring opening and the percentage of modifications in RNA by DEPC were validated using cleavage reaction and NMR. The cleavage reaction was performed on DEPC treated and untreated hairpin RNA as follows: 50 μL 1 M Tris, pH 8.0, and 25 μL of 0.4 M NaBH_4_ (prepared fresh) and incubated on ice for 30 min in the dark followed by ethanol precipitation. The pellet was resuspended in 100 μL of 1 M aniline-acetate buffer (Aniline and Glacial Acetic acid, prepared fresh) and incubated at 60°C for 15 min in the dark followed by ethanol precipitation. The resulting fragments of RNA were end-labeled by ^32^P-ATP using T4 polynucleotide kinase and run on 15 % denaturing PAGE.

NMR: The abundance of adenosine in the DEPC treated and control hairpin RNAs sequence was quantified by proton NMR spectrometry using a Bruker BioSpin NEO600 spectrometer equipped with a cryogenic probe operating at 298 K for all measurements. We have standard pulse sequence noesygppr1d with a delay of 60 seconds to allow complete relaxation of nuclei between individual scans. Assignment of proton scans was confirmed by literature and by supplemental HSQC scans.

Nucleotide modification analysis: We compared the (+)DEPC and (−)DEPC data sets and derived the final DEPC modification for adenosine using Tombo(13) method model_sample_compare with --fishers-method-context 3. The modification status of a nucleotide was assigned in Tombo by comparing the current signal distribution between (+)DEPC and (−)DEPC samples (**Fig. 1B, C**). Next, the Tombo statistic was extracted for each read and each base using custom scripts (https://git.app.uib.no/valenlab/sms-seq-pipeline). Due to Tombo working on the consensus level (average of all reads per position) we had to identify a threshold for the Tombo statistic which will allow us to work at the read level. We identified the false discovery rate (FDR) as a function of threshold using a synthetic RNA, which has adenine bases in a large number of different 5-kmer configurations (**Supplementary Table 1**). This was achieved by detecting modifications on control replicate 1 against control replicate 2 in the pipeline - and evaluating FDR for different threshold values. As the most fitting selection of a threshold is arbitrary when its relation to FDR is linear, we decided on a threshold of 0.2 (equivalent to an FDR of 15%) (**Supplementary Fig. 1)**. To evaluate it further we checked that we can still separate modified adenosine from the unmodified at the consensus level when using our threshold on individual reads (ROC curve **Fig. 1D**), resulting in AUC values ∼0.95 for different modification rates. Furthermore, we calculated FDR for the unmodified ligated RNA part of the (+)DEPC hairpin (**Fig. 1E**) to be 0.16 for the cutoff of 0.2 on the Tombo statistic (**Supplementary Fig. 2A, B**). This indicates that the variance between different replicates is not influencing the FDR greatly and the Tombo method is stable. The modification rate for the hairpin was calculated to be around 31%. When accounting for 15% of false positives in the data we estimate the real modification rate to be around 26 %, similarly to the NMR result (24.7% ± 4.27%) (**Fig. 1F**). Next, we focused on evaluating the quality of calling open-closed regions using modified-unmodified states. To achieve that we calculated ROC curves for the highly stable hairpin construct, with a well-defined loop region (**Supplementary Fig. 2**). For determining the accuracy of modification calling on the hairpin (ROC curves) we excluded reads with <4 and >18 modifications, which we deemed to be under- and over-modified, respectively (**Supplementary Fig. 2C**). Additionally, Tombo does not output any statistics for the 10 last bases of the read, and due to the --fishers-method-context set to 3 we lose an additional three bases of the read.

### Determination of dependencies

Adjusted mutual information is a variation of mutual information correcting for agreement due to chance (see **Eq. 1-2**, and ‘adjusted_mutual_info_score’ function in scikit-learn 0.22.1)(21) which can be calculated for pairs of nucleotides to see how informative knowing the modification state of one base is on predicting the state of the other base. Due to the high dependency of bases in close proximity, we normalize the AMI score between two bases by scaling AMI for each distance using z-score (dAMI). Therefore when multiple distances have high dAMI a cluster will become visible on the grid, indicating the mutual information.

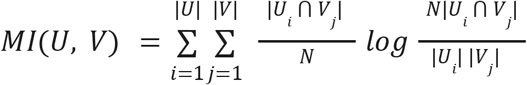

**Equation 1**. Mutual information score (MI) is calculated between two positions U and V which are represented by vectors of modified/unmodified states (corresponding to 1 and 0) for each of the reads that span both positions U and V. We required at least 10 reads between positions U and V for calculations.

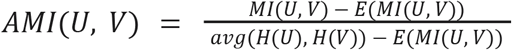

**Equation 2**. Adjusted mutual information score (AMI) normalizes MI to account for chance.

### Deciphering RNA structural heterogeneity

The hairpin RNA and snoRNAs were folded using the ViennaRNA Package (22). To determine hairpin heterogeneity, all reads were folded *in silico* with modifications used as hard constraints (not allowed to basepair). The number of reads resulting in the same structure was counted. Yeast non-coding RNAs were folded – based on sequence minimum free energy (MFE), maximum expected accuracy (MEA), and centroid – and reads were clustered to them using the binary distance metric, i.e., the proportion of reads in which *both* of the two positions are modified/open relative to the number of reads where *at least one* is modified/open. The structures of the riboswitches were obtained from previous studies (11, 23).

## Results

High-throughput RNA structure probing commonly uses reagents that react with nucleotides to interrogate the chemical accessibility of a region. The position of chemical modification is then detected by reverse transcription drop-off and/or mutation (1). Among RNA structure probing reagents 2-methylnicotinic acid imidazolide (NAI) and dimethyl sulfate (DMS) have been used for both *in vivo* and *in vitro* RNA structure determination (16, 24), whereas diethyl pyrocarbonate (DEPC) has been used to map RNA structures *in vitro* (15, 19, 20). NAI acylates the 2’OH, DMS alkylates the N-7 position of guanosine and N-3 position of cytidine (16, 24), and DEPC carbethoxylates unpaired adenosine at the N-7 position (12). To determine whether we could detect NAI, DMS and DEPC-modified bases using the Oxford Nanopore MinION sequencer we extracted *E. coli* 16S rRNA and probed it with the three reagents. The sequence quality and read length were drastically lower for NAI and DMS treated RNA compared to DEPC probed RNA (**Supplementary Fig. 3**) similar to previous reports for NAI and DMS (9, 10). The current signal in the sequencing pore is affected by 5-mer sequences occupying the reading head at a time. Thus, we assembled and cloned a 5-mer construct from oligos containing adenine bases in a diverse range of neighboring nucleotide sequence contexts. We synthesized RNA from this construct to probe using DEPC (see **Methods**). Using this construct we compared the output from sequencing unmodified RNA molecules to RNA modified with DEPC at different concentrations (1%, 5%, 10%, and 5% for denatured RNA). The comparison was performed using Tombo which can detect significant differences in the ionic current running through pores when presented with a modified versus unmodified base (**Fig. 1B**)(13). This comparison demonstrated that modified bases could be distinguished from unmodified bases (**Fig. 1C**) at high accuracy (AUC values ∼0.95, **Fig. 1D**). This is consistent with other studies that demonstrated chemical reagent-induced RNA modifications can be detected using Oxford nanopore sequencing (9, 10).

To assess how well this approach could capture structural information, we synthesized and probed a hairpin RNA with a known, stable structure (**Supplementary Fig. 2A**). We first determined DEPC modification through footprinting analysis where cleavage patterns indicate modifications in a single-stranded region of the hairpin (**Fig. 1E**). We then quantified the modification level of the hairpin RNA treated with DEPC using proton NMR. This showed a decrease of adenosine integral by 24.7% ± 4.27% (**Fig. 1F**). We sequenced hairpin RNA treated with DEPC and compared these with control samples obtaining 3093 and 4687 full-length reads, respectively. All hairpins were ligated to a longer unmodified molecule and the transition from the unmodified to the modified hairpin was readily distinguishable as a deviation from the expected output from the Oxford Nanopore MinION sequencer (**Supplementary Fig. 2B**) confirming that modifications are readily detectable using nanopore sequencing.

Recently, a nanopore sequencing-based method demonstrated consensus calling of RNA structure (9, 10). Consensus calling calculates an average across all reads and therefore produces consensus structures, but RNA can potentially form a heterogeneous ensemble of structures (25). We, therefore, sought to call modifications at the level of individual bases on single molecules as this makes it possible to detect heterogeneous structures formed by the same sequence, modification dependencies across the molecule, and RNA breathing – the transient opening of RNA structure. Using Tombo we set a threshold for detecting individual bases on individual reads based on the false discovery rate (FDR). The FDR was calculated by predicting modifications in the untreated sequences ligated to the hairpins and reads from the unmodified 5-mer construct (see **Methods**). We selected a threshold of 0.2, corresponding to a false discovery rate of ∼15%, which when applied to the hairpin (**Fig. 1G**), resulted in an average of 7.43 modified bases over full-length reads (**Supplementary Fig. 2C**). This corresponds to a modification rate of 26% and is consistent with the modification rate obtained from NMR. We observed a significant enrichment of modified bases in the open region versus the stem of the hairpin (**Fig. 1H, I**) and obtained near-perfect consensus classification across all reads when considering open versus closed (stem) adenines (**Fig. 1J, Supplementary Fig. 4, 5A**). More interestingly, individual bases on individual reads achieved a sensitivity of 0.50 and specificity of 0.74 (**Fig. 1J**). However, since the NMR results indicate that most accessible bases are not modified in every read, the number of false negatives is highly inflated and the sensitivity is therefore underestimated. False positives in the inaccessible regions come, at least partially, due to the nature of nanopore sequencing. Some k-mers are more sensitive than others to be detected as modified and modifications of neighboring nucleotides can influence the signal (**Supplementary Fig. 5A**). However, some modifications within stems are likely to represent real, low-frequency conformations where these bases are transiently exposed and modified (“RNA breathing”) (3). To explore this, we folded individual reads with the ViennaRNA Package (22) using hard constraints prohibiting modified bases from being part of a stem. Reassuringly, the most abundant structure was the expected hairpin structure indicating that 45.1% of reads have no modification conflicting with this structure. These were however heavily modified in the expected open regions (**Fig. 1K** inset, **Supplementary Fig. 5B**), showing that the hairpin structure is indeed the most favorable conformation. Most alternative structures are only from a single read, and potentially represent noise in the calling of modifications (**Fig. 1K**). However, some structures were observed more frequently and showed bases modified in the hairpin stem and forming larger loop regions (**Fig. 1K**, inset). The frequency of these more abundant structures was correlated with the minimum free energy (MFE) (**Fig. 1K**) indicating that these could represent transient structures. We explored this heterogeneity using an enzymatic cleavage assay (**Supplementary Fig. 5C**) which revealed the presence of low-abundance open regions across the stem. Together these observations are consistent with the heterogeneity predicted from SMS-seq. This suggests that SMS-seq can measure structures beyond the consensus structure and reveal conformational heterogeneity or RNA breathing.

Methods based on enzymatic cleavage or termination during reverse transcription can only capture independent measurements from a single location. Unlike these, SMS-seq can modify and record numerous exposed bases per molecule. SMS-seq is therefore capable of capturing dependencies of open regions across individual molecules. We first assessed the extent to which we could obtain information on the dependencies by calculating the adjusted mutual information (AMI) between all pairs of bases (see **Methods**). Mutual information measures how much information the state of one base informs of the state of another base. Since calling the modification state of a base using nanopore sequencing is influenced by its neighboring bases, we scaled the AMI according to the distance between the pair of bases considered. Using z-score normalization (see **Methods**), this provided us with a distance-normalized AMI (dAMI). Applying this metric to the hairpin revealed that bases that are part of the same structural unit share a high dAMI (**Fig. 2A**). In particular, we observed that the two base-paired regions forming the stem of the hairpin had a modification state that depends on each other (**Fig. 2A, black rectangle**) and to a lesser extent the bases in the loop (**Fig. 2A, black triangle**). While the dAMI can infer dependencies it does not distinguish between open and closed states, but only estimates the extent of information captured by one position in relation to the other. We, therefore, also calculated to what extent bases are co-observed in the same state. Specifically, we calculated *co-openness*, the probability that one base is modified given that the other is modified; and *co-agreement*, the fraction of reads where the pair share the same state (**Fig. 2B**). These metrics highlighted the open and stem regions, respectively. Together this demonstrated that SMS-seq can capture structural co-dependencies in stable structures.

**Fig. 2.**
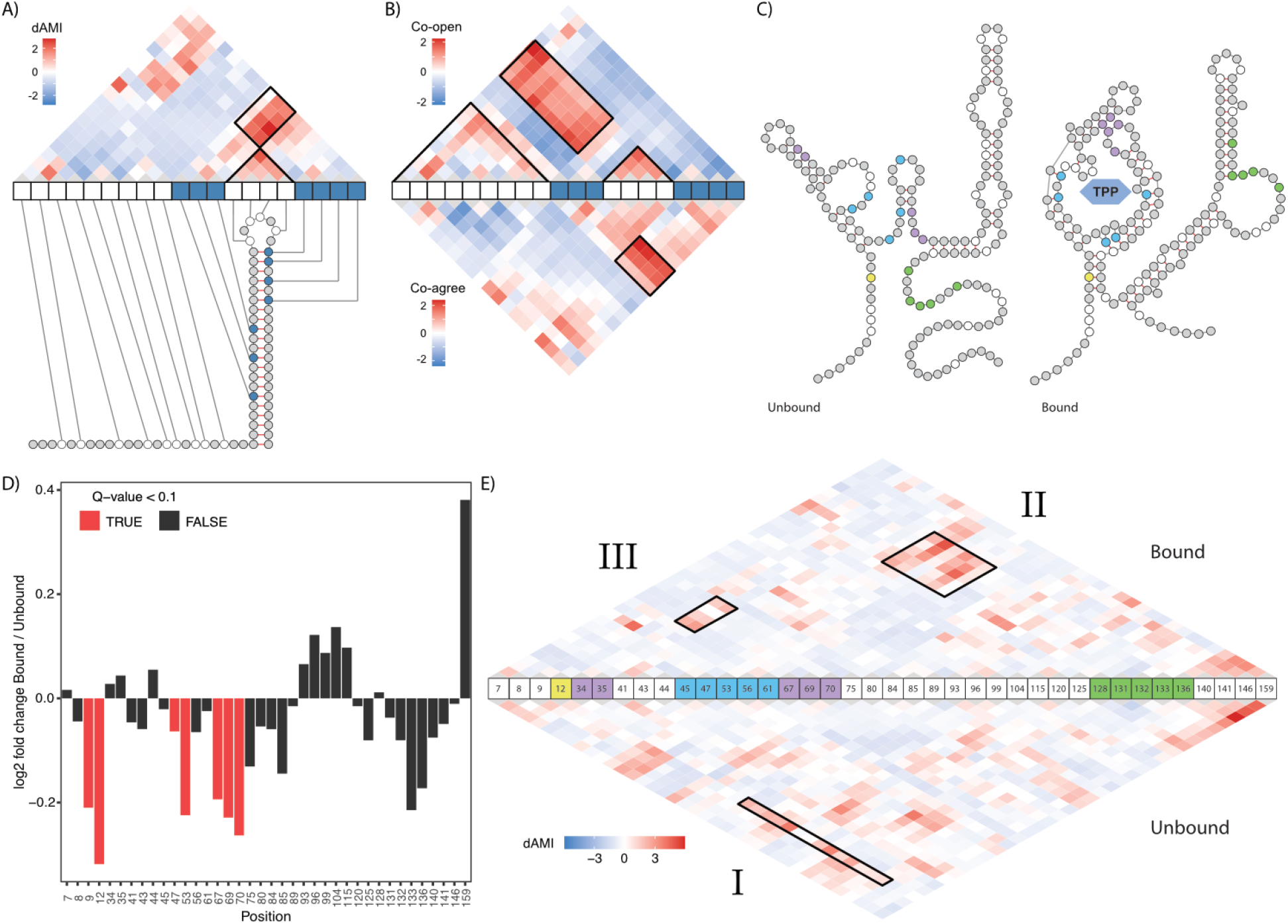
SMS-seq reveals structural dependencies in RNA. **A)** dAMI dependencies for pairs of bases in the synthetic hairpin construct. **B)** Z-score normalized co-openness and co-agreement scores for the synthetic hairpin construct. **C)** The predicted secondary structure obtained from a previous study(23) for the TPP riboswitch in unbound (left) and bound (right) configurations. Domains are indicated in colors: first stem (yellow), aptamer domain (blue), tertiary interaction of loop-stem (purple), and ribosome binding site (green). Only adenosine residues are colored. **D)** Log2 fold change in accessibility between bound and unbound states of TPP. Only adenosine residues are depicted. Red bars indicate significant changes (FDR adjusted Cochran–Mantel–Haenszel test q < 0.1) in the ligand-binding domain. **E)** dAMI dependencies between pairs of bases for the bound (upper) and unbound (lower) state of the TPP riboswitch. Domains are colored according to subfigure C. Three black rectangle labels as I, II, and III show some of the differences between bound and unbound TPP.

We next asked whether SMS-seq could capture the dynamics and dependencies of structured riboswitches. We probed two riboswitches in their ligand-bound and unbound states: thiamine pyrophosphate (TPP (23), **Fig. 2C**) and *F. nucleatum* Flavin mononucleotide (FMN (11), **Supplementary Fig. 6A**). As expected, both riboswitches became overall less accessible upon ligand-binding (**Fig. 2D, Supplementary Fig. 6B**). Comparing between bound and unbound states of TPP revealed that, in particular, the aptamer domain (positions 11–88) exhibited significant differences in accessibility (**Fig. 2D**). The structural dependencies predicted by dAMI further showed that the aptamer domain and those that encompass the ribosome binding site (positions 125-146) are interdependent. This observation is consistent with the reduced access for ribosomes and translational inhibition upon ligand binding as previously observed (26). The high accessibility of the P1 helix, consisting of the pairing of base 11-14 with 85-88 (represented by position 12, **Fig. 2C**, yellow dot, and **Fig. 2D**) in the unbound state showed a strong decrease in accessibility in the bound state indicating its role in conformational change upon ligand recognition. Our dAMI analysis further supported this by showing the role of the first stem (represented by position 12) on the overall architecture of TPP (**Fig. 2E**, a diagonal line extending from 12) including the ribosome binding site (**Fig. 2E, box I**). The dependency of the expression domain on the conformational change of the TPP binding pocket in the presence of TPP is also evident (**Fig. 2E, box II**). Interestingly, dAMI also revealed that SMS-seq is capable of capturing the tertiary interactions between the two helices (**Fig. 2E, box III**), which has been demonstrated in mutational profiling (RING-MaP) and high-resolution TPP structure mapping (3, 23), where the interaction between nucleotides in the L5 loop (bases from 67 to 72) and the P3 helix (a stem structure formed by base pairing of nucleotide 21-27 and 33-38) enables the formation of the ligand-binding pocket. Although RING-Map and SMS-seq agree on loop-helix interaction, they are not completely consistent in their predictions (**Supplementary Fig. 7**). For instance, RING-MaP suggests RNA interaction groups of L5 with P2 (base pairing between 15-18 and 48-51) and L5 with P4 (base pairing between 56-60 and 80-83), which might not be true since the hinge-like movement of helix P5 relative to P4 (27) may not allow such an interaction (**Supplementary Fig. 7**). Similar observations can be made for the FMN where SMS-seq captures the extensive dependencies (stem-loop and loop-loop) of the two wings of FMN (**Supplementary Fig. 6C, D**). Together, these results demonstrate that SMS-seq can capture secondary and tertiary dependencies and the conformational changes of riboswitches upon ligand binding.

To investigate the structural heterogeneity of RNA molecules in solution, we folded the FMN riboswitch in different concentrations of magnesium ion (28, 29). Similar to a previous report (29) SMSseq revealed different modification patterns, suggesting heterogeneous conformations of FMN (**Supplementary Fig. 6E**).

Encouraged by the results on synthetic RNA structures, we set out to investigate the structure of non-coding RNAs and generated non-poly(A) RNA libraries from yeast (*Saccharomyces cerevisiae*) and folded these *in vitro* in the presence and absence of DEPC. We first aligned the reads to *in silico* folded structures and to known structures from RNACentral (30). For structures such as the small nucleolar RNA (snoRNA) snR37 and snR17a where we had many reads, we observed overall agreement between reads and their reported or energetically favorable structures (**Fig. 3A, B** showing snR37). To investigate whether reads could represent structural heterogeneity we attempted to cluster reads to multiple alternative conformations predicted by ViennaRNA: the minimum free energy, maximum expected accuracy, and centroid structure for snR17a. We observed that reads clustered preferentially to these specific structures (**Fig. 3C**) and rejected randomly shuffled versions of these (t-test p < 2.2e-16, see Methods). We similarly obtained thousands of reads for rRNAs showing modification patterns in agreement with sub-structures of this long molecule (**Supplementary Fig. 8)**. Together these findings highlight that SMS-seq can capture multiple structures of an RNA.

**Fig. 3.**
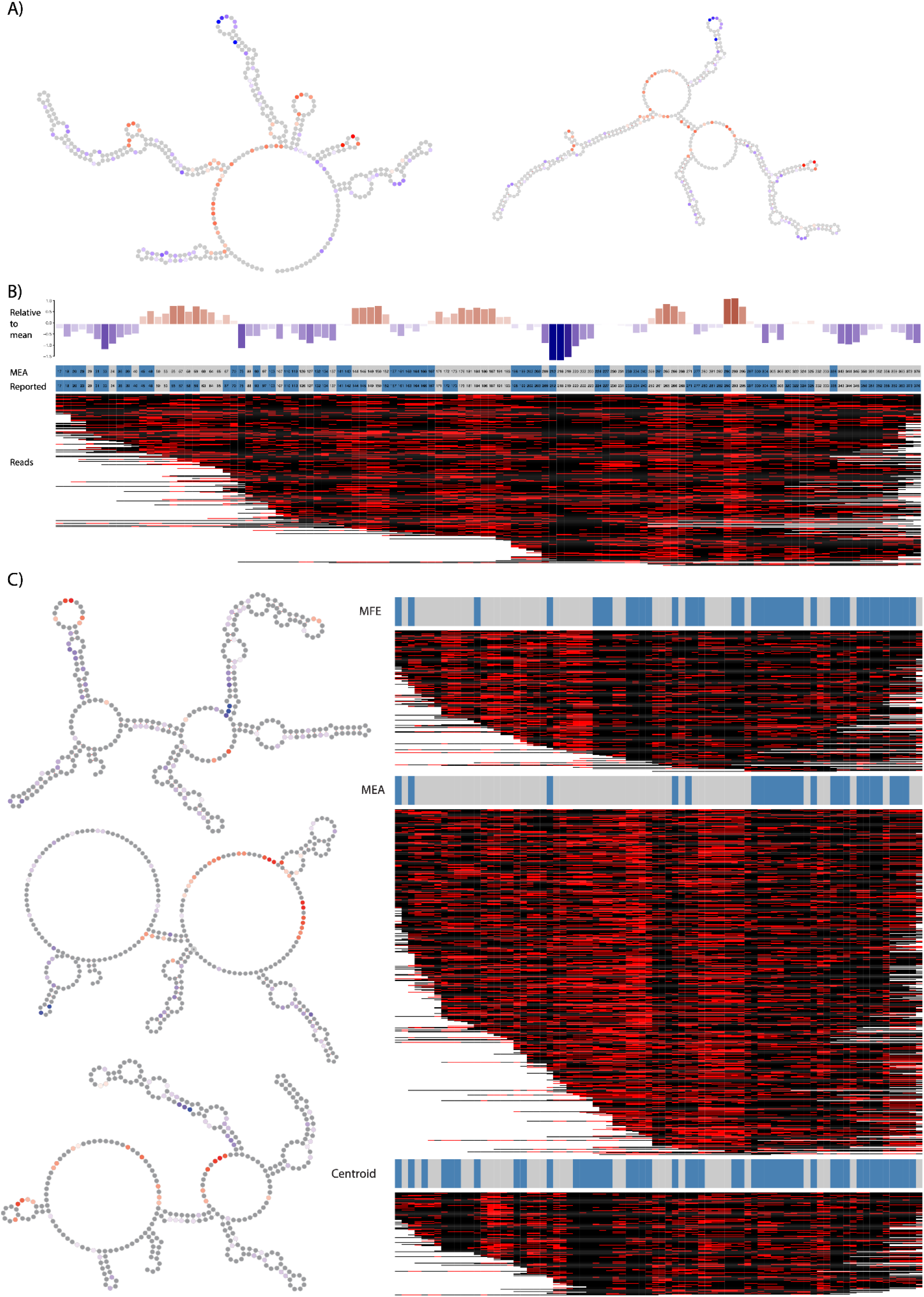
Structural probing of non-coding RNA. **A)** Two structures of snR37, *in silico*-folded maximum expected accuracy (MEA, left) and structure from RNACentral(35) (right). Bases are colored according to their accessibility relative to the mean accessibility for this molecule. **B)** Accessibility per position relative to mean (upper panel), Structure by position for MEA and Reported structures (mid panel) where stem (blue) and open (gray), and all reads mapping to snR37 with bases colored according to whether they are modified (red) or not (black), (lower panel). **C)** Reads mapping to snR17a clustered by their similarity to three *in silico* predicted structures. On the left, structures represent folding based on minimum free energy (MFE), maximum expected accuracy (MAE), and centroid. On the right, nanopore reads are clustered towards each of the predicted structures (the red in the heatmap and the gray bars on the top of it mark the open regions). To assess the concordance of read modifications to *in silico* predicted structures we clustered reads against these and randomly permuted structures ten-thousand times. This demonstrated that reads overwhelmingly agreed with predicted structures compared to permuted (t-test p-value < 2.2e-16).

We finally expanded our SMS-seq approach to endogenous mRNA and isolated poly(A) RNA from yeast and probed these *in vitro* using DEPC. Average modification rates were highly reproducible across replicates (**Supplementary Fig. 9**) and similar modification patterns were observed along the mRNAs (**Supplementary Fig. 10**). We compared these mRNA-selected libraries with libraries based on enzyme-based PARS-assisted folding (31), and observed a significant difference in modification rates between nucleotides classified by PARS as open and closed (**Supplementary Fig. 11A**). We could also observe known structural elements in mRNAs (**Supplementary Fig. 11B**) in line with a previous study (32). Globally, our approach was able to confirm known properties of mRNA structure showing an increase in modifications upstream of translation start sites and translation stop sites consistent with these regions being open (31) (**Fig. 4A**). Furthermore, 5’UTRs overall showed a higher modification rate and higher accessibility compared to CDSs, while 3’UTRs were overall less accessible (**Fig. 4B**) At the global level the average modification rate among mRNAs varies considerably (**Fig. 4C**). Some genes like HAC1 are predicted to form extensive base pairing consistent with reports that HAC1 form elaborate structures through long-range base pairing for intron splicing (33). Other transcripts appear to be overall unstructured. To demonstrate the utility of SMS-seq we investigated more specific regions to study their structural features. For instance, by splitting translation termination sites into the 3 different stop codons (UAA, UGA, UAG), SMS-seq revealed that modification rates decreased at the beginning of 3’UTR for UGA stop codons specifically (**Fig. 4D**). SMS-seq also predicted more accessible regions around several polyadenylation signal motifs (**Fig. 4E**). Together these observations show that SMS-seq can be used to probe structural features of the transcriptome.

**Fig. 4.**
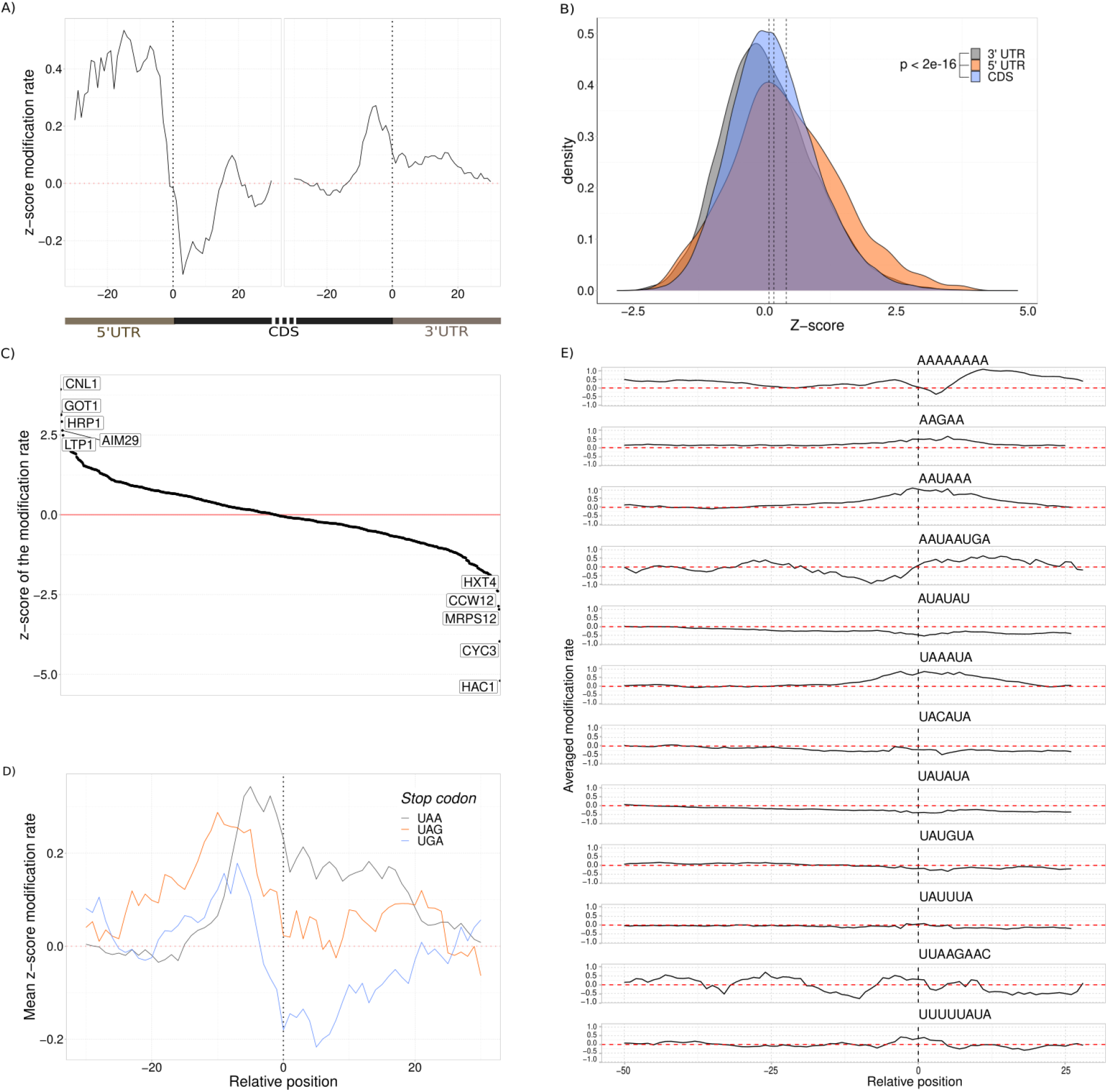
mRNA accessibility and features. A) Metaplot of the genes over translation initiation site and translation termination sites. Each gene is individually z-score normalized and then averaged across all genes. Positions with a z-score below 0 are less accessible than average. B) Comparison of CDS and UTR modification rate. Significant differences were detected using Student’s t-test. C) Accessibility around translation termination sites grouped by stop codons. D) Genes ranked by their average modification rate show a range of accessibilities. The five lowest and highest are labeled by their gene name. E) Polyadenylation signal sequences of yeast mRNA and their modification rate 50 nt upstream and 20 nts downstream. The zero position represents the start of the sequence motif.

## Discussion

We here introduced a method for single-molecule probing of RNA structures, SMS-seq, that does not rely on amplification or size selection, and does not need to be normalized based on position in the molecule – common sources of biases in previous methods (1, 34). Unlike many methods, SMS-seq does not rely on reverse transcription thus avoiding another source of bias. We have shown that SMS-seq can detect transient and stable RNA structures, secondary and tertiary structural elements, structural dependencies of bases, and mRNA structural features.

Since SMS-seq is a single-molecule method it obtains structural information from individual RNA molecules and can therefore be used to uncover heterogeneity and the dynamic nature of RNA structures in different conditions. A previous method, RING-MaP, could capture within molecule correlations of modified bases at the consensus level (3, 23); however, they are incapable of capturing longer RNA-RNA interactions. Other clustering-based algorithms like DREEM (6) and DRACO (7) are able to detect heterogeneity of RNA structure but suffer from experimental artifacts of mutational profiling and short-read sequences when used to determine larger RNA structures.

Previously, nanopore direct RNA sequencing was demonstrated in the determination of structures of pre-miRNA clusters and mRNA isoforms (9, 10). While these results revealed consensus structures, SMS-seq probes multiple bases in each individual molecule and uses this to investigate the interdependency of these positions and the overall structure. While obtaining these dependencies requires substantial coverage, we expect that with the rapid improvement to modification-calling algorithms, the sensitivity of our approach will increase and can eventually be used as a constraint for RNA structure prediction algorithms.

An important aspect of SMS-seq and other nanopore-based structure probing methods is that the sensitivity of the method to detect long-range interaction largely depends on a high modification rate combined with high integrity of the RNA. RNA structure probing using SHAPE reagents (2-methylnicotinic acid imidazolide (NAI), 1-methyl-7-nitroisatoic anhydride (1M7), and N-methylisatoic anhydride (NMIA)), dimethyl sulfate (DMS) and nanopore sequencing resulted in low quality, truncated and unmappable sequences (9, 10). In contrast, DEPC resulted in multiple modifications per read and a higher read integrity at the expense of only being applicable for *in vitro* experiments. However, as a method, SMS-seq can be effectively adapted for RNA-structure *in vivo* probing assays when *in vivo* probing agents obtain comparable high-modification, high-integrity reads.

Another shortcoming of DEPC is its lack of ability to modify other bases than adenosine. Although a new probing reagent that could target all 4 nucleotides, acetylimidazole, has been synthesized to use in nanopore-based structure determination, it is shown that at the higher concentrations necessary to obtain extensive modification rate the quality and fraction of full-length reads were reduced (9). Further improvements in RNA probing reagents without compromising the RNA integrity are expected to advance RNA structure determination.

Building on our and other long-read-based approaches with increased accuracy and structure prediction, we expect full-length single-molecule structural probing of native RNA by direct RNA sequencing will eventually become a standard tool in predicting secondary structure and inferring structural dependencies and tertiary interactions.

### Data and materials availability

The sequencing files are available on the European Nucleotide Archive website, under accession code PRJEB36658. Scripts used to process the data are available on https://git.app.uib.no/valenlab/sms-seq.

## Acknowledgments

We would like to acknowledge Ruth Brenk (University of Bergen) for providing FMN riboswitch RNA. We also acknowledge our lab members for detailed and constructive comments that helped to improve this study. The study was funded by Norwegian Research Council (project number: 250049) and Trond Mohn Foundation.

## Author contributions

T.T.B. conceived the study, performed the principal experiments, analyzed part of the results, and wrote the manuscript. K.L. analyzed the data, wrote part of the manuscript. M.J. performed NMR analysis. K.J. performed part of the experiments. A.N. contributed for data analysis. EV conceived the study, supervised the study, analyzed part of the results, and wrote the manuscript. All authors read and approved the manuscript.

