## Supplementary for "Single molecule structure sequencing reveals RNA structural dependencies, breathing and ensembles"

### Supplementary Materials

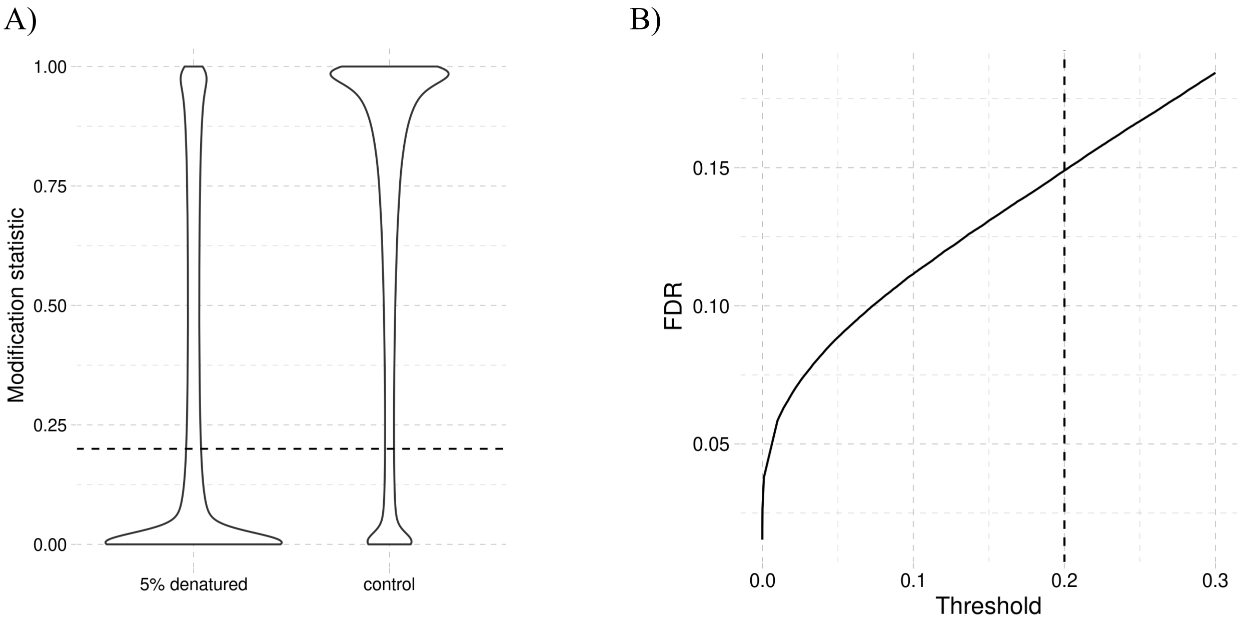


**Supplementary Figure 1. K-mer construct and statistical thresholds**. K-mer construct that contains 5-mer contexts that allow checking the modification level of adenine. **A)** A denatured RNA was treated with 5 % DEPC and untreated control. The selected modification statistic threshold of 0.2 is marked with a dashed line. **B)** The false discovery rate (FDR) for the selected threshold (0.2) resulted in an FDR of ~0.15.

A)

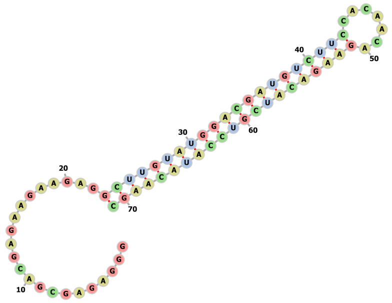

B)

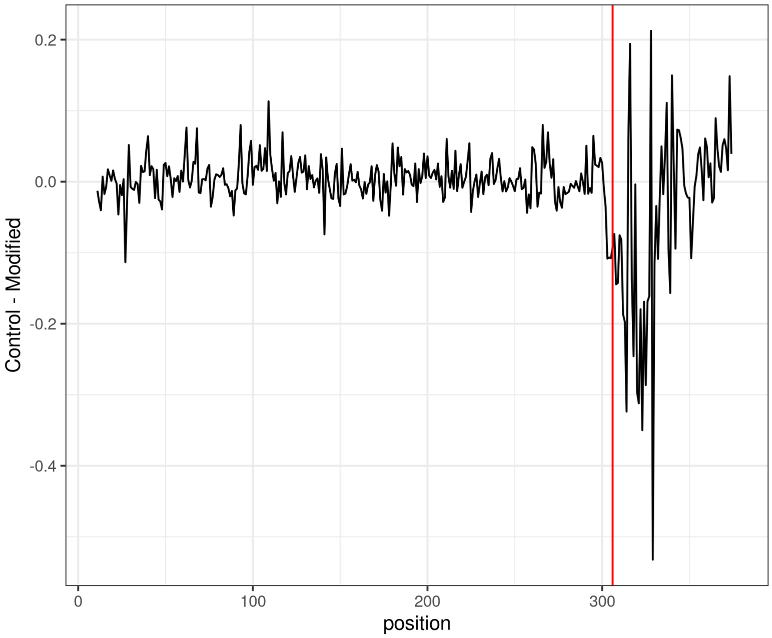
Control - Modified
0.2
0.0
-0.2
-0.4
0
100
200
300
position

C)

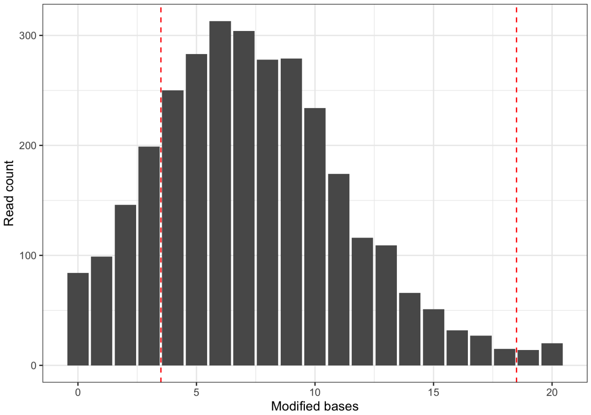
Read count
300
200
100
0
0
5
10
15
20
Modified bases

**Supplementary Figure 2. Structural probing of a synthetic hairpin RNA.**

**A)** The structure of the synthetic hairpin used for evaluation. The shown secondary structure is a minimum free energy structure folded with Vienna RNAFold(1). **B)** The mean difference (in pico ampere) in Nanopore current signal of an RNA-ligated hairpin between reads in the DEPC-modified and control libraries. In both these libraries, an unmodified RNA (position 0-306, left of the red line) is ligated to a hairpin (position > 306, right of the red line). The hairpins are modified in the DEPC-modified library and unmodified in the control library. **C)** Number of reads with a given modification frequency. The number of full-length reads spanning the hairpin (y-axis) split by the number of modified adenines (x-axis) in the read. Reads with <4 or >18 modifications corresponding to almost completely modified or almost unmodified reads, respectively, are excluded from the accuracy estimation (see Methods).

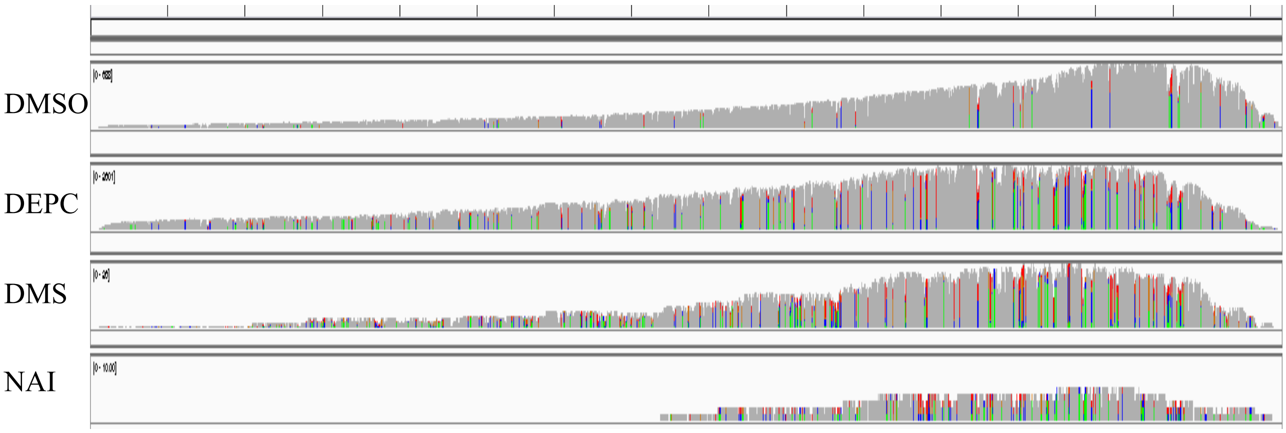


**Supplementary Figure 3.** **Testing of different RNA structure probing reagents**. *E. coli* 16S rRNA was treated with DEPC, DMS, NAI, and untreated control. The IGV read coverage plot on the full-length 16S rRNA depicts NAI and DMS treated RNA had short and low-quality sequences. Whereas DEPC treatment had yielded a full-length read comparable with the control. The y-axis is the number of reads covering a given position and the x-axis is the base position along 16S rRNA. Each bar represents a nucleotide. The gray bars represent an exact match and the colored bars indicate at least 10 % of the reads have mismatches to the reference sequence.

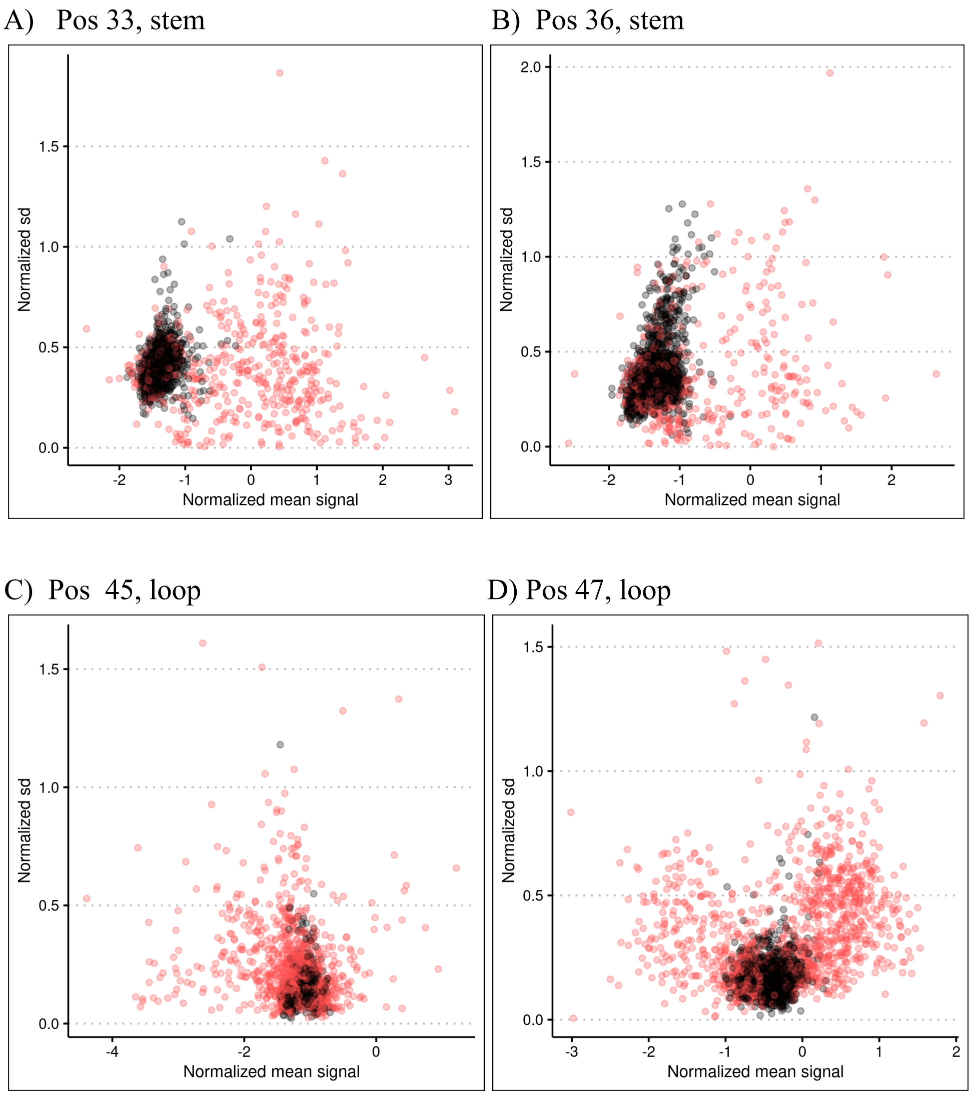


**Supplementary Figure 4. Raw data for synthetic hairpin RNA.**

Normalized signal and standard deviation carry information to distinguish modified (red) and unmodified (black) bases and in turn to recognize which regions are open (loop) and closed (stem). The x-axis displays a normalized mean raw nanopore signal, while the y-axis displays normalized standard deviation of the raw nanopore signal.

A)

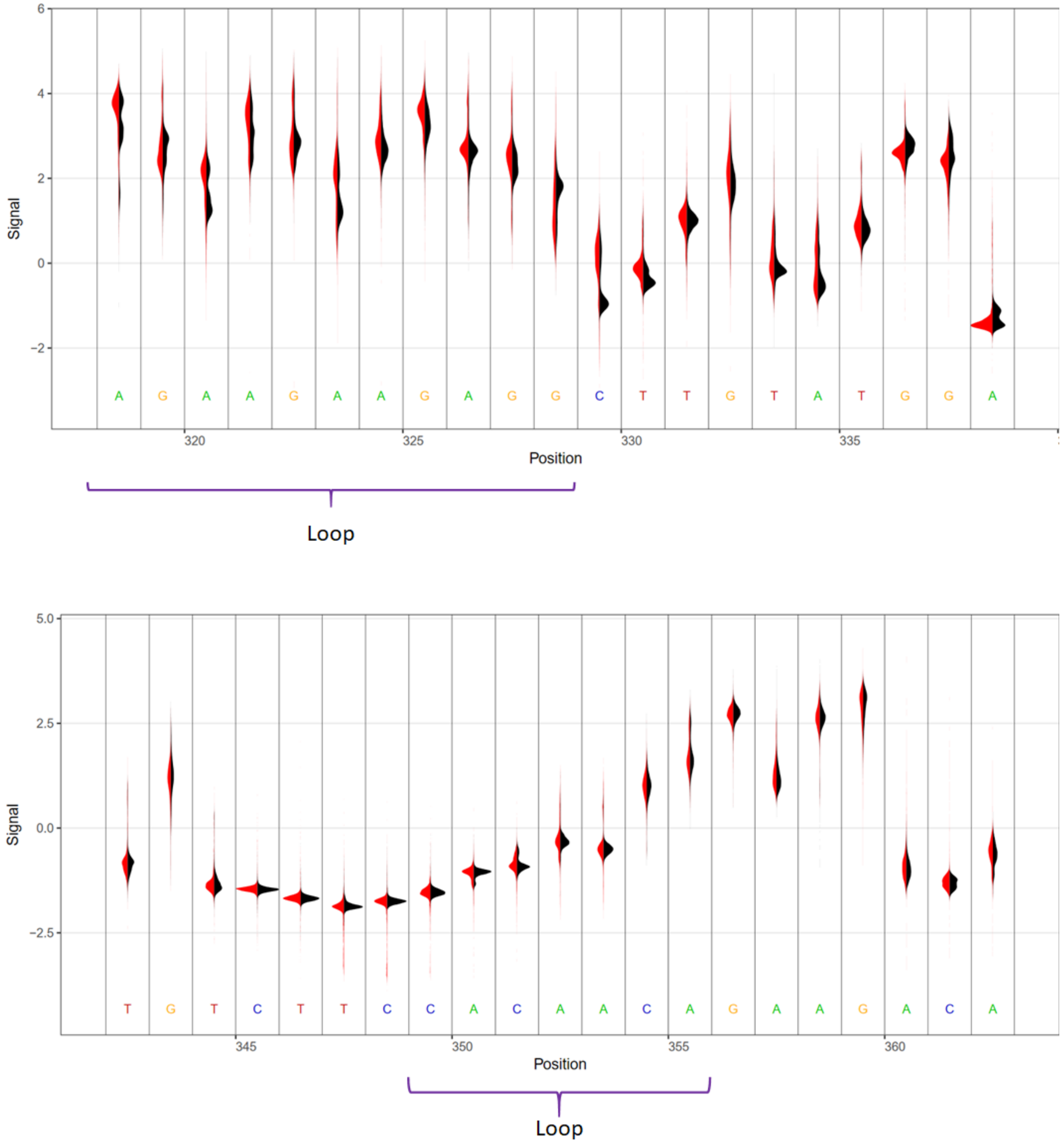
Signal
6
4
2
0
−2
A G A A G A A G A G G C T T G T A T G G A
320
325
330
335
Position
Loop
Signal
5.0
2.5
0.0
−2.5
T G T C T T C C A C A A C A G A A G A C A
345
350
355
360
Position
Loop

B)

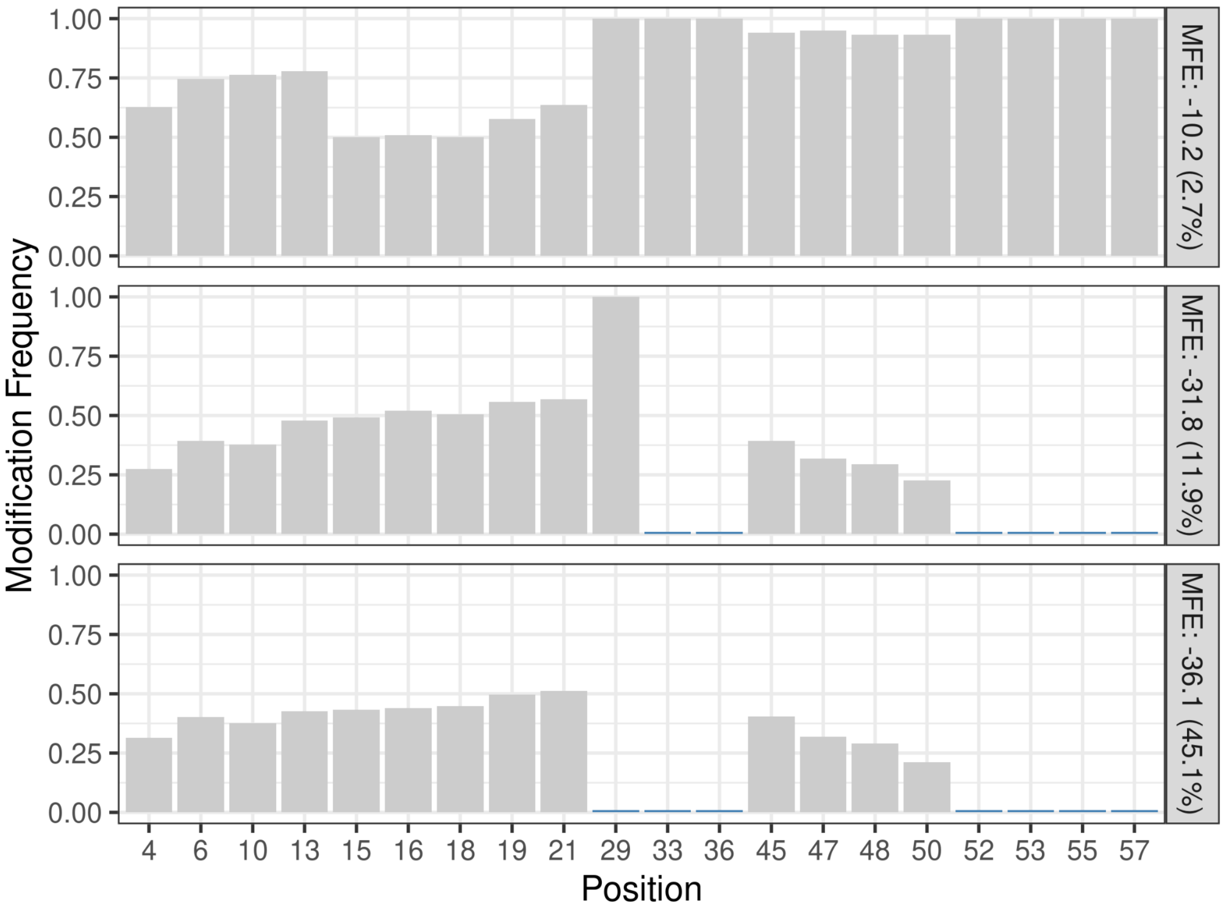

Modification Frequency
1.00
0.75
0.50
0.25
0.00
MFE: -10.2 (2.7%)
MFE: -31.8 (11.9%)
MFE: -36.1 (45.1%)
4
6
10
13
15
16
18
19
21
29
33
36
45
47
48
50
52
53
55
57
Position

C)

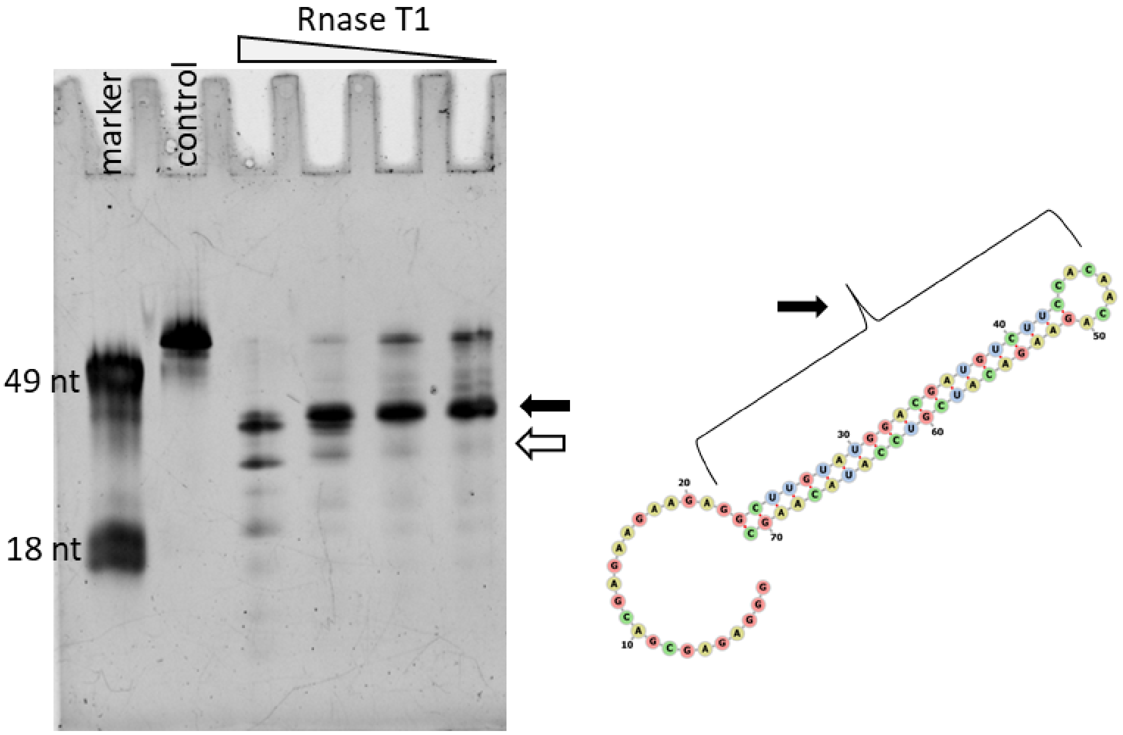


**Supplementary Figure 5. Nanopore raw current signal and alignment for the synthetic hairpin and enzymatic assay.**

A) Distribution of Nanopore resquiggled current signal from control (black) and DEPC treated (red) hairpin RNA. Predicted accessible regions are labeled as “loop”. B) Modification frequency per base over synthetic hairpin for selected three RNA structures with different minimum free energy (MFE). C) RNase T1, which cuts after G, enzymatic cleavage of hairpin RNA shows the predominant structure (black arrow) and alternative structures with an opening of the stem (white arrow).

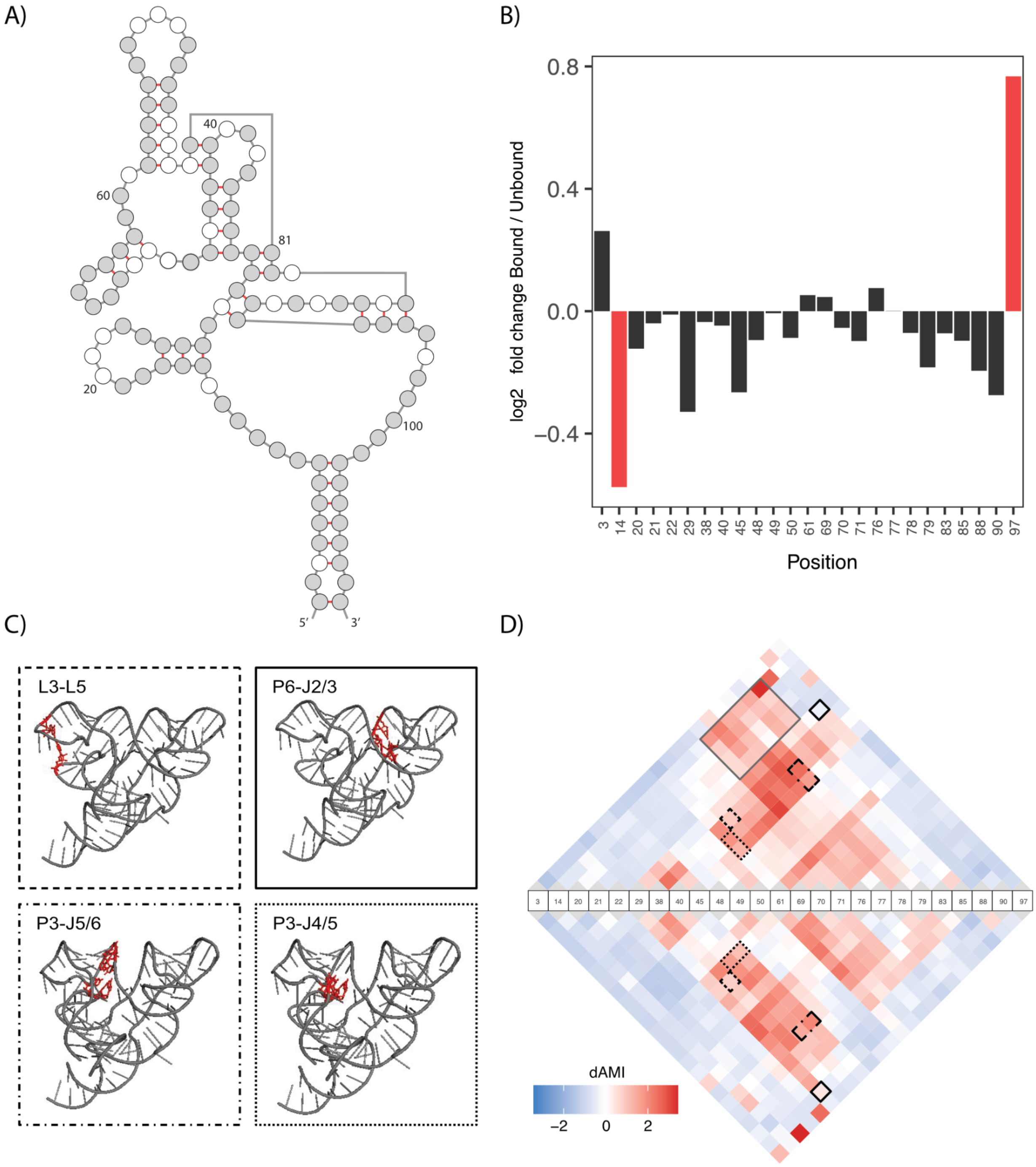
A)
40
60
81
20
100
5′
3′
B)
0.8
0.4
0.0
−0.4
log2 fold change Bound / Unbound
3 14 20 21 22 29 38 40 45 48 49 50 61 69 70 71 76 77 78 79 83 85 88 90 97
Position
C)
L3-L5
P6-J2/3
P3-J5/6
P3-J4/5
D)
3 14 20 21 22 29 38 40 45 48 49 50 61 69 70 71 76 77 78 79 83 85 88 90 97
dAMI
−2
0
2

E)

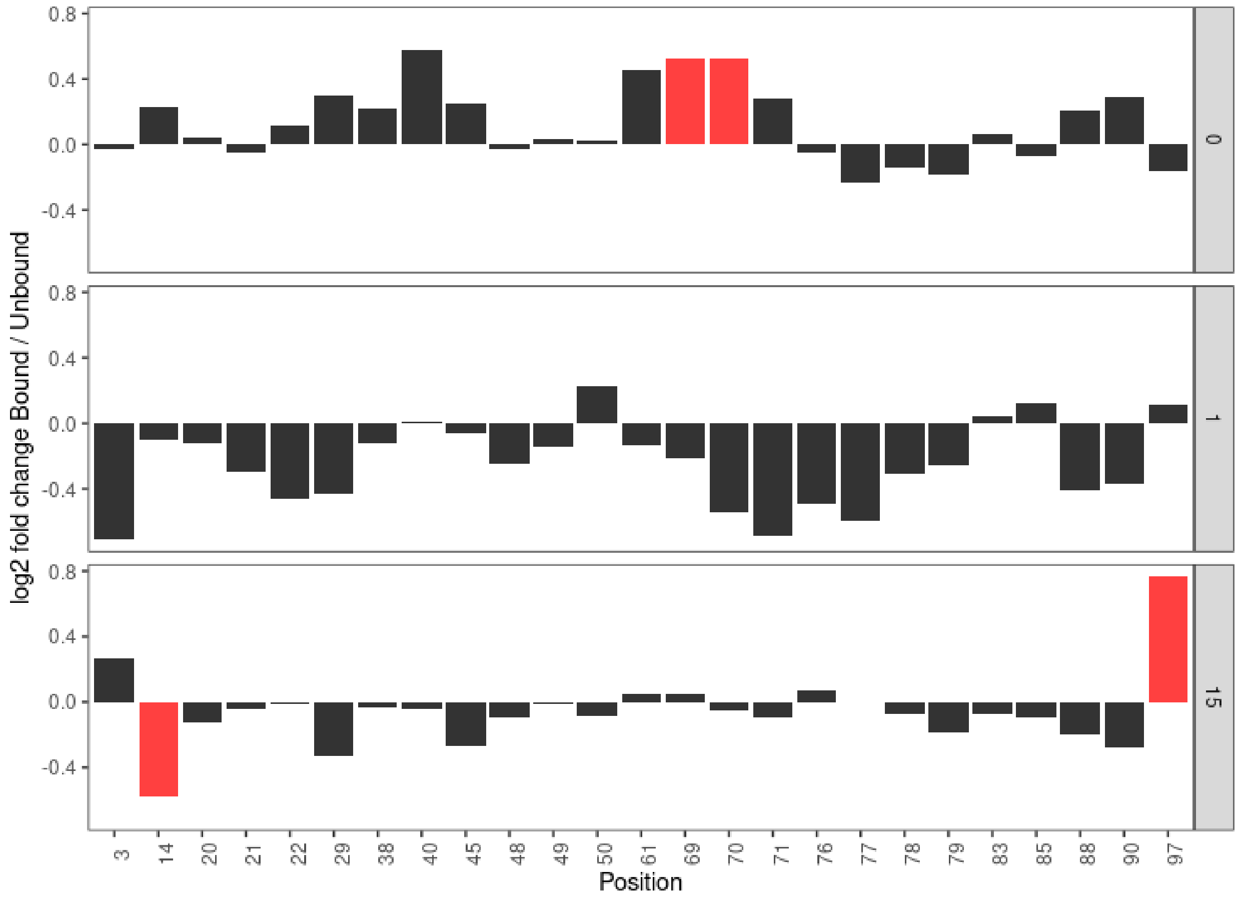


**Supplementary Figure 6.** **FMN riboswitch.**

**A)** The FMN riboswitch RNA structure. **B)** Log2 fold change in accessibility between bound and unbound states of FMN. Red bars indicate significant change. Adding the ligand-induced conformational changes reveals the accessibility of some of the nucleotides at the ligand-binding pockets. **C)** FMN riboswitch RNA in a stick-and-ribbon representation (PDB ID: 3F2Q(2)) and tertiary interactions are coloured red using PyMOL (v 1.5)(3). L, P, and J stand for loop, stem, and junction, respectively. **D)** dAMI dependencies between pairs of bases for the bound (upper) and unbound (lower) state of the FMN riboswitch, which reveals inter-domain dependencies and tertiary interactions where the ligand essentially lock-up the riboswitch with more dependencies among different domains (the top gray big box). Nucleotides reorientation upon ligand binding induces a change in the dependencies among different domains. Some of the tertiary interactions that have been described using X-ray crystallography(2) are depicted in the dependencies plot with different line styles boxes designated in the panels (C). **E)** Log2 fold change in accessibility between bound and unbound states of FMN in 0 mM, 1 mM, and 15 mM MgCl2. Red bars indicate significant change. The different concentration of Mg2+ induced heterogeneous conformations threby the accessibility of some of the nucleotides (4).

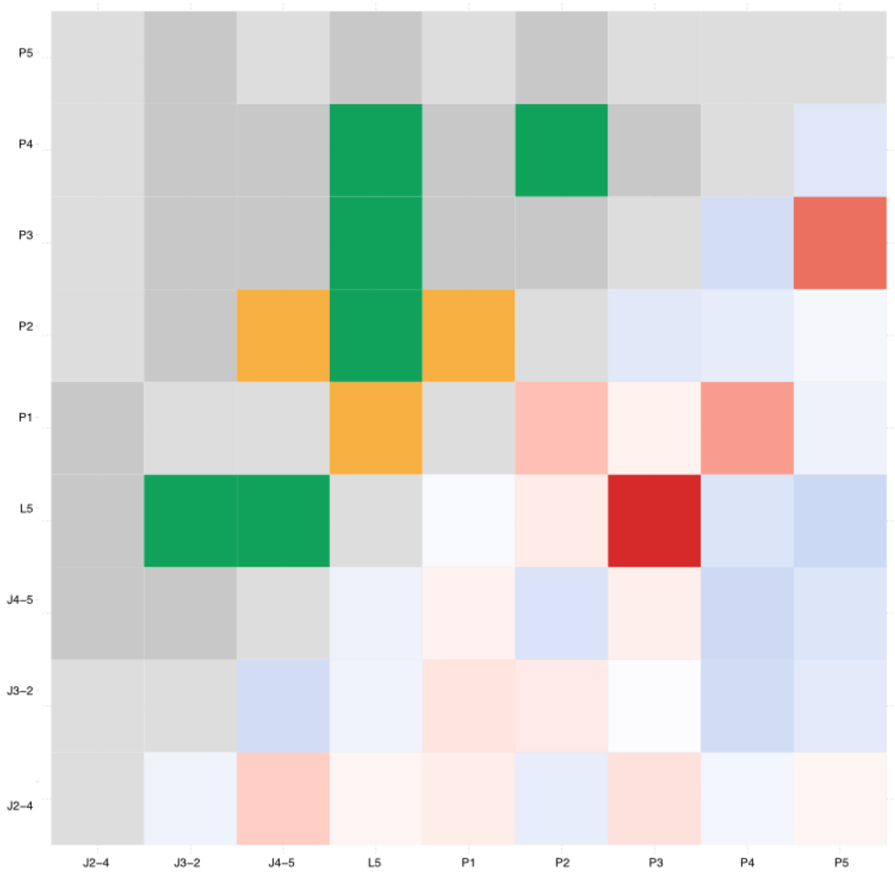


**Supplementary Figure 7. Comparison between RING-MaP data and dAMI metric of TPP.** RING-MaP correlation (left side) and SMS-seq dAMI (right side). RNA interaction groups among regions in TPP. J, P, and L refer to junction, stem and loop regions, respectively. dAMI is sensitive in picking up for example the known TPP structural interaction between nucleotides in P3 and L5 region in the presence of ligand(5), while not highlighting false positives of RNA interaction groups of L5 with P2 and L5 with P4.

A)

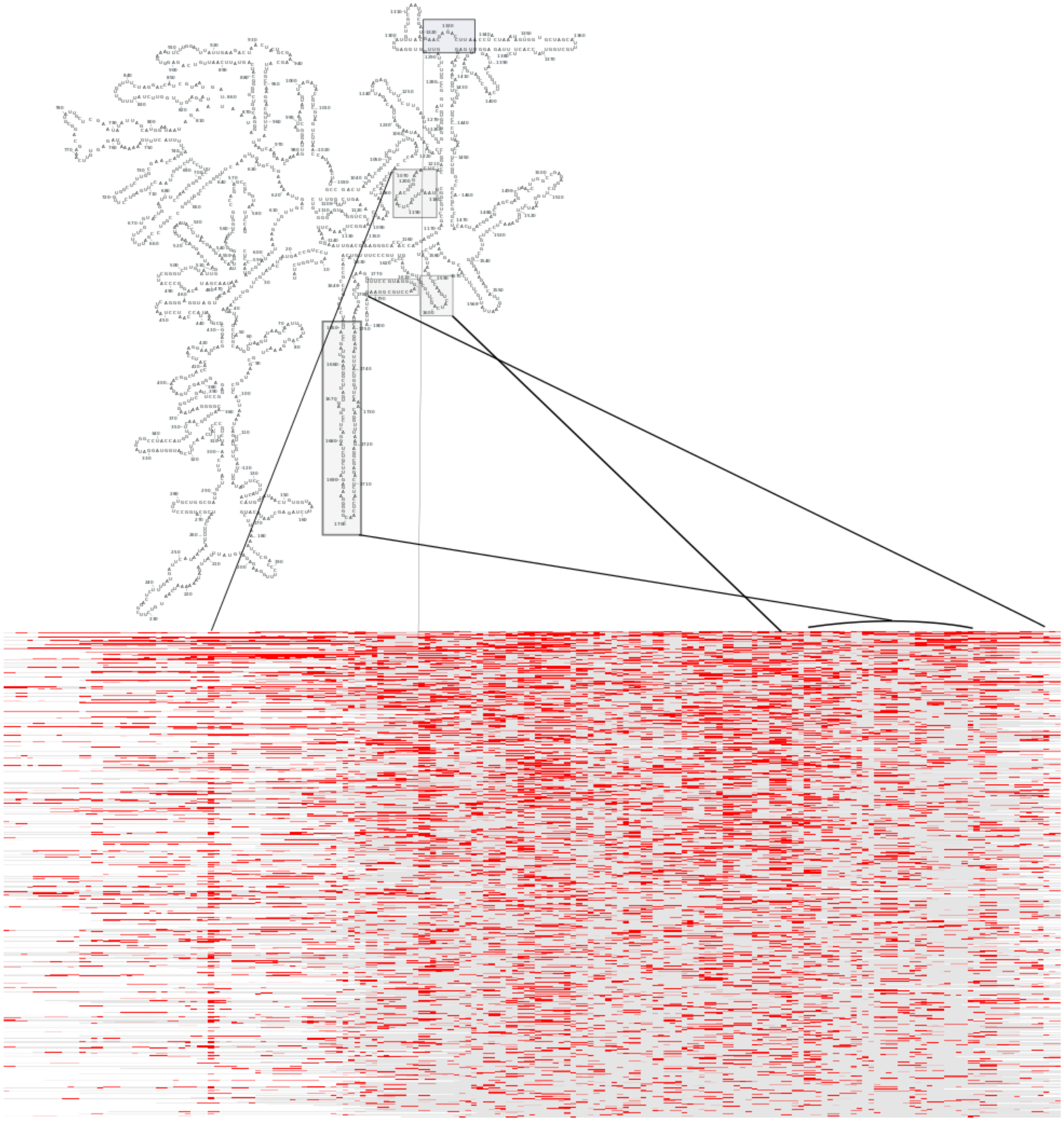

B)

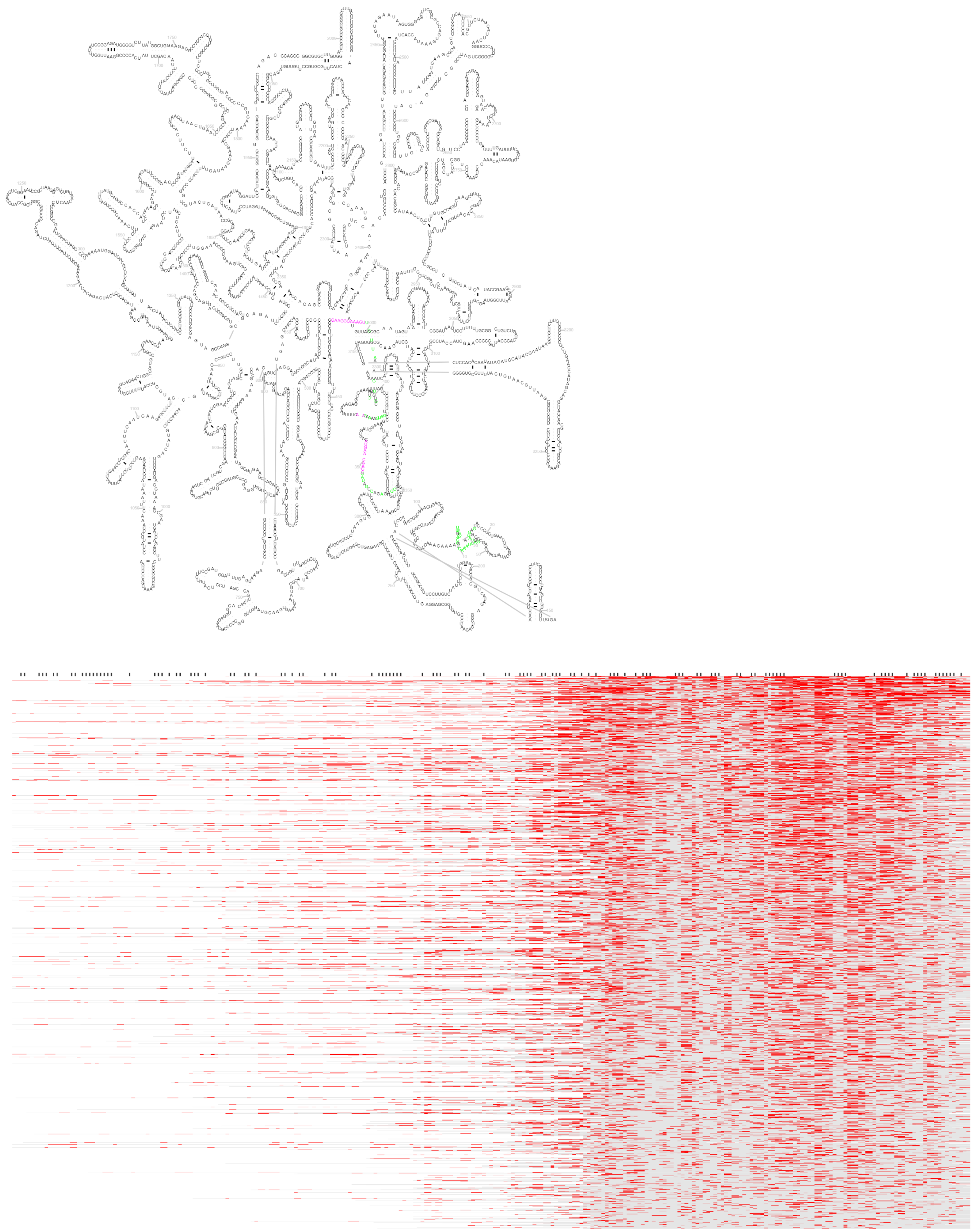

**C)**

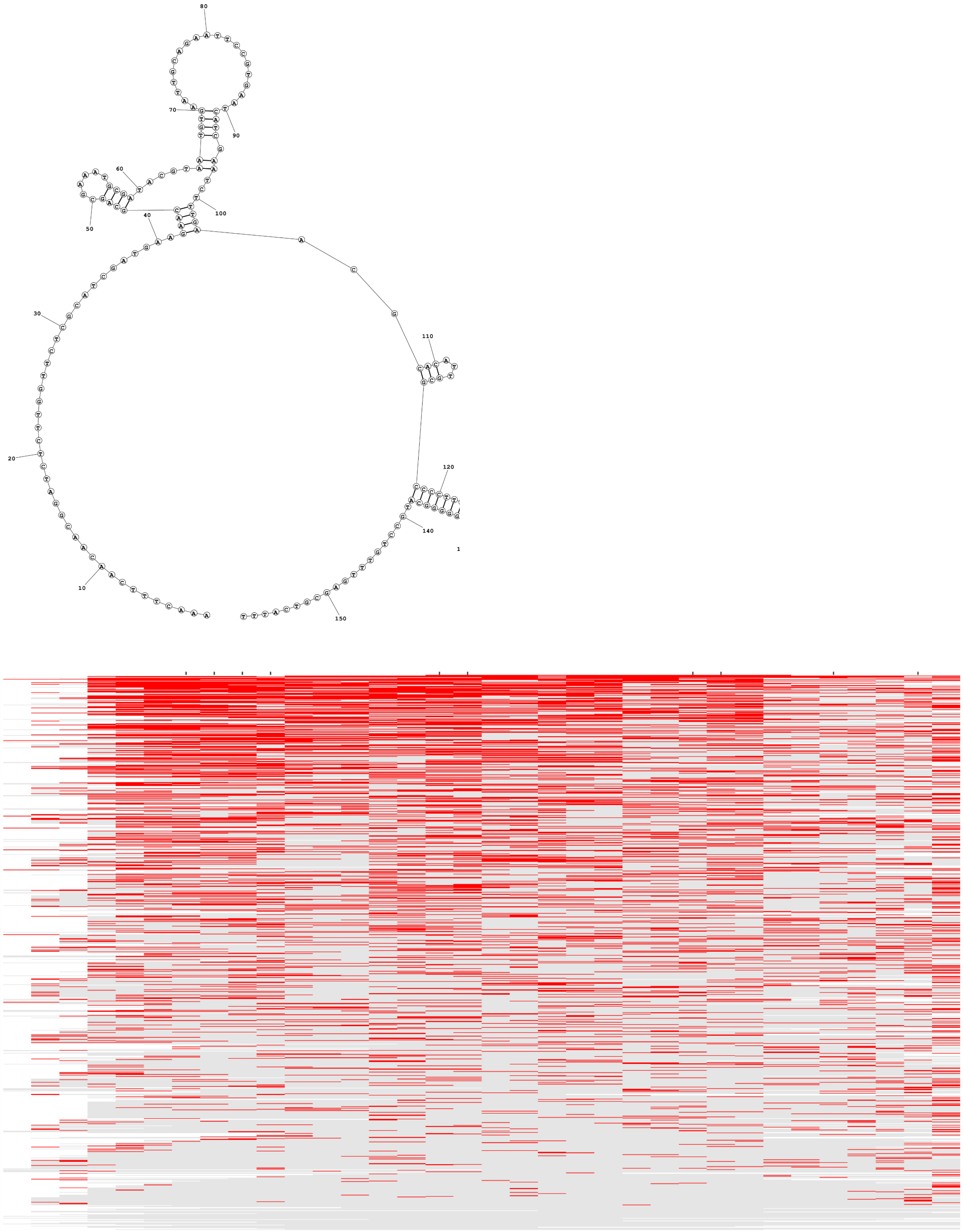

D)

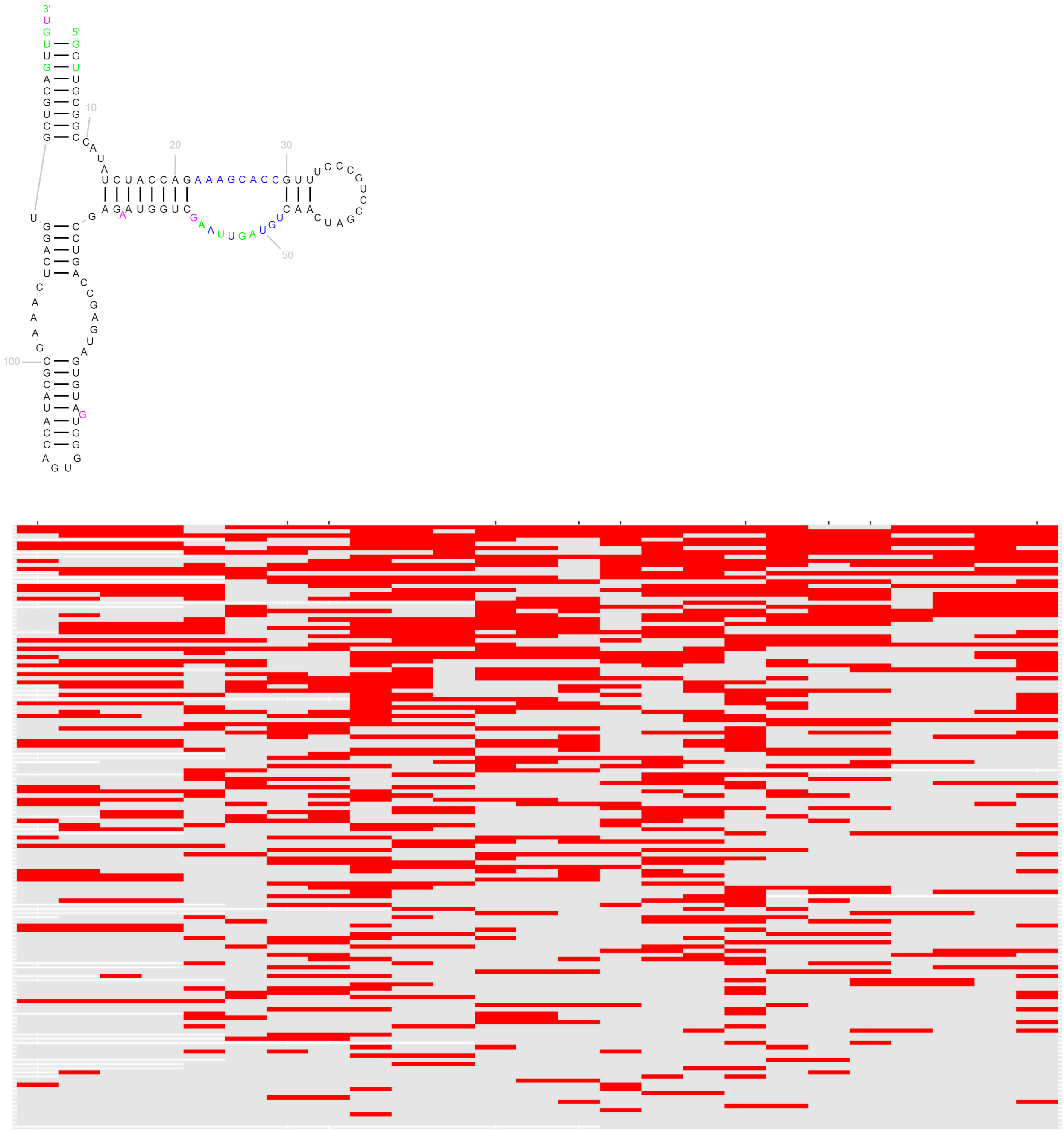


**Supplementary Figure 8.** **Structural probing of yeast rRNA.**

Phylogeny-based secondary structure generated using RiboVision (top) and the modification level (bottom) of A) the 3′ ends of 18S rRNA, B) the 3′ ends of 25S rRNA, C) 5.8S rRNA, and D) 5S rRNA. Red and gray on the heatmap indicate modified and unmodified positions, respectively.

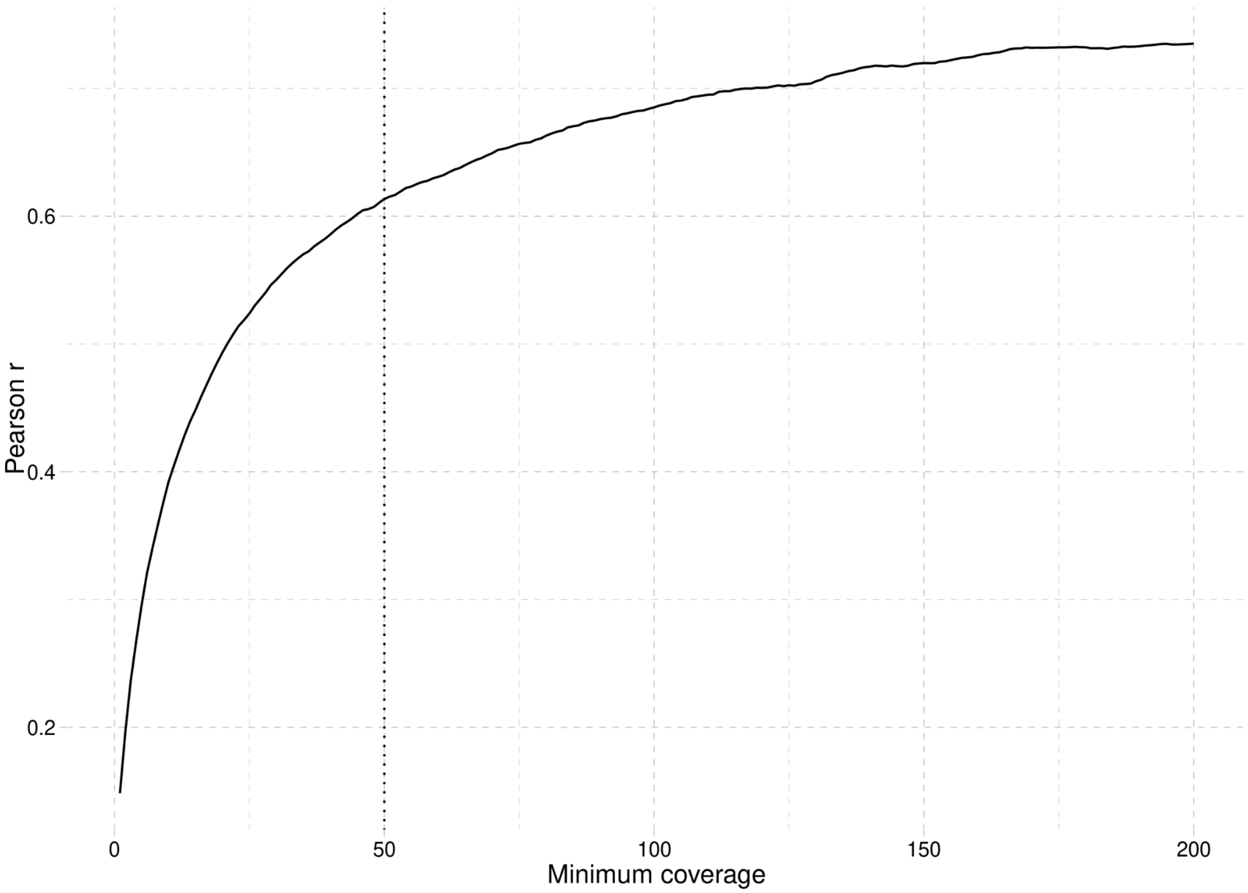


**Supplementary Figure 9. Relation between coverage and correlation for two yeast replicates.** Correlations here are calculated based on z-score - modification rate normalized per gene per replicate. We selected 50 as minimum coverage for being included in further analysis, which results in above 0.6 correlations between replicates for modification rate.

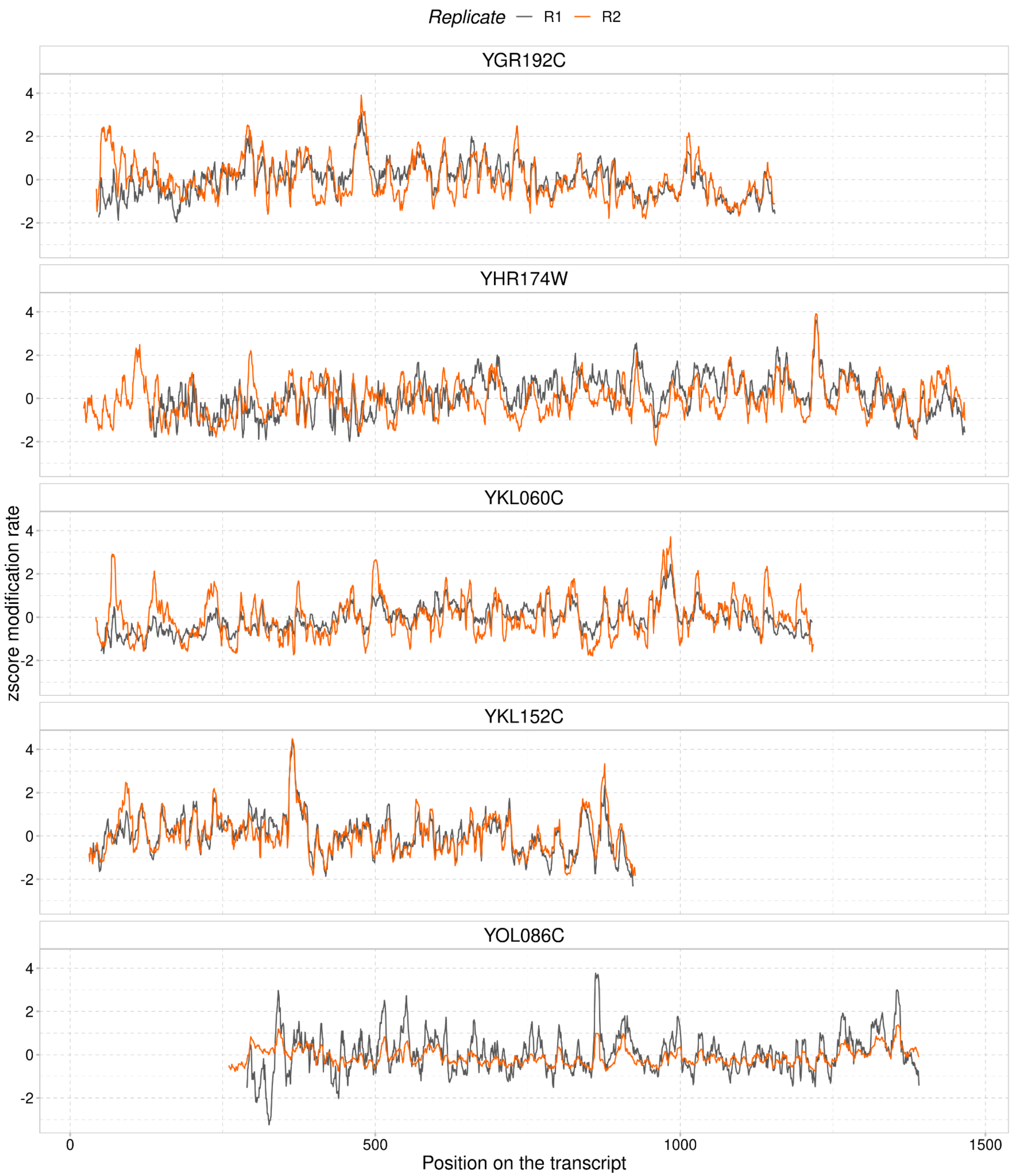


**Supplementary Figure 10. Modification rate along highly expressed mRNAs.** Z-score of the modification rate correlates well within individual genes between replicates. The pattern of low/high z-score seems to be very well preserved.

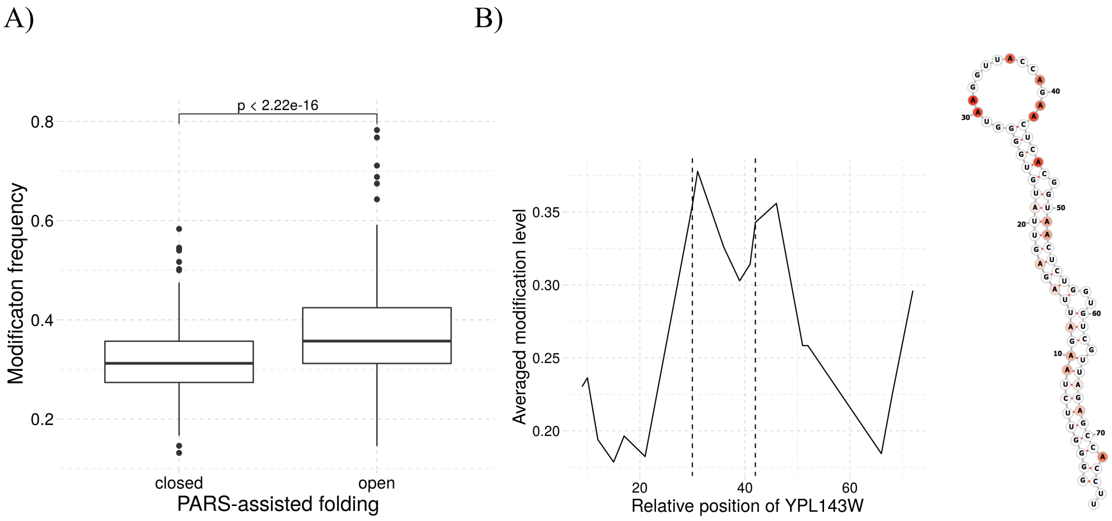


**Supplementary Figure 11. Comparison of SMS-seq to PARS-assisted folding and SMS-seq results for a known mRNA structure.** A) The SMS-seq modification frequency for regions that are classified as either open or closed according to PARS-assisted folding in yeast mRNAs (6). B) Modification signal of a known hairpin structure in mRNA. The structure is from RPL33A (YPL143W) mRNA (chromosome XVI:282824-282900) and agrees with previous *in vitro* DMS structural probing (7). SMS-seq modification signal is color-coded proportional to modification level and plotted onto the RNAfold structure prediction (right). Vertical broken lines define the loop region.

**Supplementary Table 1.** RTA adapters, TPP oligos and RNA sequences ligated at the 5′ end of the hairpin and riboswitches.

| RTA adapters | |
|---|---|
| Barcode name | Sequence |
| BC1 Oligo A | 5'-/5Phos/CCTCCCCTAAAAACGAGCCGCATTTGCGTAGTAGGTTC-3' |
| BC1 Oligo B | 5'-GAGGCGAGCGGTCAATTTTCGCAAATGCGGCTCGTTTTTAGGGGAGGTTTTTTTTTT-3' |
| BC2 Oligo A | 5'-/5Phos/CCTCGTCGGTTCTAGGCATCGCGTATGCTAGTAGGTTC-3' |
| BC2 Oligo B | 5'-GAGGCGAGCGGTCAATTTTGCATACGCGATGCCTAGAACCGACGAGGTTTTTTTTTT-3' |
| BC3 Oligo A | 5'-/5Phos/CCTCCCACTTTCACACGCACTAACCAGGTAGTAGGTTC-3' |
| BC3 Oligo B | 5'-GAGGCGAGCGGTCAATTTTCCTGGTTAGTGCGTGTGAAAGTGGGAGGTTTTTTTTTT-3' |
| BC4 Oligo A | 5'-/5Phos/CCTCCTTCAGAAGAGGGTCGCTTCTACCTAGTAGGTTC-3' |
| BC4 Oligo B | 5'-GAGGCGAGCGGTCAATTTTGGTAGAAGCGACCCTCTTCTGAAGGAGGTTTTTTTTTT-3' |
| TPP Oligos | |
| Oligo-1 | 5'-TAATACGACTCACTATAGGGCCAAACGACTCGGGGTGCCCTTCTGCGTGAAGGCTGAGAAATACCCGTATCACCTGATCTGGATAATGCCAGCGTAGGGAA-3' |
| Oligo-2 | 5'-AGCTTGCGCTGAACCCAGCAGGTCGACTTGCATAGTTTGCTCCTGCCATAACGTGAAGAAGCAATGACCTGGTGGTCCGTGACTTCCCTACGCTGGCATTAT-3' |
| Sequences of ligated RNAs | |
| Ligated RNA to hairpin | 5'-GCGCUCCAUGCAAACCUGUCCUCGAAGCAUUGUAAGUCGCCACCAUGGUGAGCAAGGGCGAGGAGCUGUUCACCGGGGUGGUGCCCAUCCUGGUCGAGCUGGACGGCGACGUAAACGGCCACAAGUUCAGCGUGUCCGGCGAGGGCGAGGGCGAUGCCACCUACGGCAAGCUGACCCUGAAGUUCAUCUGCACCACCGGCAAGCUGCCCGUGCCCUGGCCCACCCUCGUGACCACCCUGACCUACGGCGUGCAGUGCUUCAGCCGCUACCCCGACCACAUGAAGCAGCACGACUUCUUCAAGUCCGCCAUGCCCGAAGGCUACGUCCAGGAGCGCACCAUCUUCUUCAAGGACGACGGCAACUACAAGACCCGCGCCGAGGAUGCCAAAAAAAAAA-3' |

| | |
|---|---|
| Ligated RNA to riboswitches | 5'-GCGCUCCAUGCAAACCUGUCCUCGAAGCAUUGUAAGUCGCCACCAUGGUGAGCAAGG GCGAGGAGCUGUUCACCGGGGUGGUGCCCAUCCUGGUCGAGCUGGACGGCGACGUAAA CGGCCACAAGUUCAGCGUGUCCGGCGAGGGCGAGGGCGAUGCCACCUACGGCAAGCUGA CCCUGAAGUUCAUCUGCACCACCGGCAAGCUGCCCGUGCCCUGGCCCACCCUCGUGACC ACCCUGACCUACGGCGUGCAGUGCUUCAGCCGCUACCCCGACCACAUGAAGCAGCACGA CUUCUUCAAGUCCGCCAUGCCCGAAGGCUACGUCCAGGAGCGCACCAUCUUCUUCAAGG ACGACGGCAACUACAAGACCCGCGCCGAGGAUGCCAA-3' |
